# Evolution of Translation Initiation Factor 2 Extensions Links Initiation to Bacterial Stress Response

**DOI:** 10.64898/2026.02.08.704698

**Authors:** Evrim Fer, Rajdeep Banerjee, Katsumi Hagino, Kaitlyn M. McGrath, Betül Kaçar

## Abstract

Terminal extensions are recurrent features in protein evolution, often linked to environmental adaptation and novel regulatory or interaction functions. Here, we combine comparative genomics, structural modeling, and functional assays to elucidate the evolutionary diversification and functional significance of one of the key proteins of all cells, translation initiation factor 2 (IF2) terminal extensions across the tree of life. Specifically, we reconstruct the first comprehensive evolutionary map of IF2 across life, analyzing ∼800 homologs and classify seven distinct structural architectures of IF2 based on extension regions. These extensions are enriched for intrinsically disordered and phase-separation–promoting residues, suggesting roles beyond the conserved catalytic core. Further, functional characterization of IF2 with varying N-terminal lengths show that loss of the N-terminal extension slows bacterial growth specifically under temperature and pH stress. Appending C-terminal extensions from different organisms to the *Escherichia coli* IF2 demonstrates a conserved role for these extensions in adaptation to temperature and anaerobiosis. Our findings establish the functional significance of IF2 terminal extensions, linking their evolutionary diversification to stress-dependent regulation of translation.

## Introduction

Protein synthesis is a core cellular process carried out by the ribosome through the sequential stages of initiation, elongation, and termination. In bacteria, translation initiation requires three initiation factors, whereas archaea and eukaryotes employ a larger and more complex set of factors^1–4^. Initiation Factor 2 (IF2) is a universally conserved translational GTPase that catalyzes essential steps of initiation, including initiator tRNA recruitment and ribosomal subunit joining^5–7^. Although the biochemical functions of IF2 are well characterized, its evolutionary diversification and potential contributions to regulation of translation across environmental conditions have not been examined^8,9^.

In bacteria, IF2 promotes initiator tRNA binding to the ribosomal P-site and facilitates subunit joining immediately prior to elongation, whereas its archaeal and eukaryotic homologs (aIF5B and eIF5B, respectively) function primarily in subunit joining ^3,5,10–14^. All IF2/IF5B homologs share a highly conserved catalytic core containing GTP- and tRNA-binding domains, yet they differ markedly in the presence, length and composition of their terminal extensions^14^. Bacterial IF2 often contains long N-terminal extensions that are dispensable for growth under laboratory conditions^15–17^, whereas archaeal IF5B lacks an N-terminal extension entirely. Conversely, C-terminal extensions are more prevalent in archaea and eukaryote homologs. In eukaryotes, these regions have been proposed to interact with eIF1A to promote ribosomal subunit joining^18–23^, but the function of corresponding extensions in archaea remains unknown. Together, these differences raise a central evolutionary problem: why distinct lineages have retained different terminal extensions on an otherwise conserved initiation factor, and what functional roles these regions serve.

Across protein evolution, terminal extensions frequently emerge as adaptive modules, often enriched in intrinsically disordered residues and capable of conferring regulatory or interaction capacities in response to environmental change^24–28^. However, despite growing appreciation of extension regions in various proteins, whether such extensions play analogous roles within translation factors has not been examined. Here, we address this gap by combining comparative genomics, structural modeling, and functional assays to reconstruct the evolutionary diversification and to assess the functional significance of IF2 terminal extensions across the tree of life. Using progressive N-terminal truncations of *Escherichia coli* IF2 and chimeric constructs bearing terminal extensions from bacterial, archaeal, and eukaryotic homologs, we assess translational performance and cellular growth across temperature, pH, and anaerobic conditions. Integrating these data with phylogenetic analyses, we identify seven distinct IF2 architectures shaped by evolutionary history. Together, our results demonstrate that lineage-specific IF2 terminal extensions evolved as modulators of translation under stress, revealing an evolutionary link between translation initiation and cellular stress adaptation.

## Results

### Evolutionary landscape and selection pressure across IF2 sequences

To reconstruct the evolutionary landscape of IF2, we analyzed 1,003 representative genomes spanning Bacteria, Archaea, and Eukaryotes filtered from an initial dataset of ∼25,000 bacterial and ∼2,700 archaeal and 1,236 eukaryotic genomes (**Fig. 1a, see Materials and Methods**). We identified 809 high-confidence IF2 homologs (563 bacterial, 234 archaeal, 12 eukaryotic) that together capture the full domain diversity in both the N-terminal and C-terminal regions of IF2 across the tree of life.

**Fig 1.**
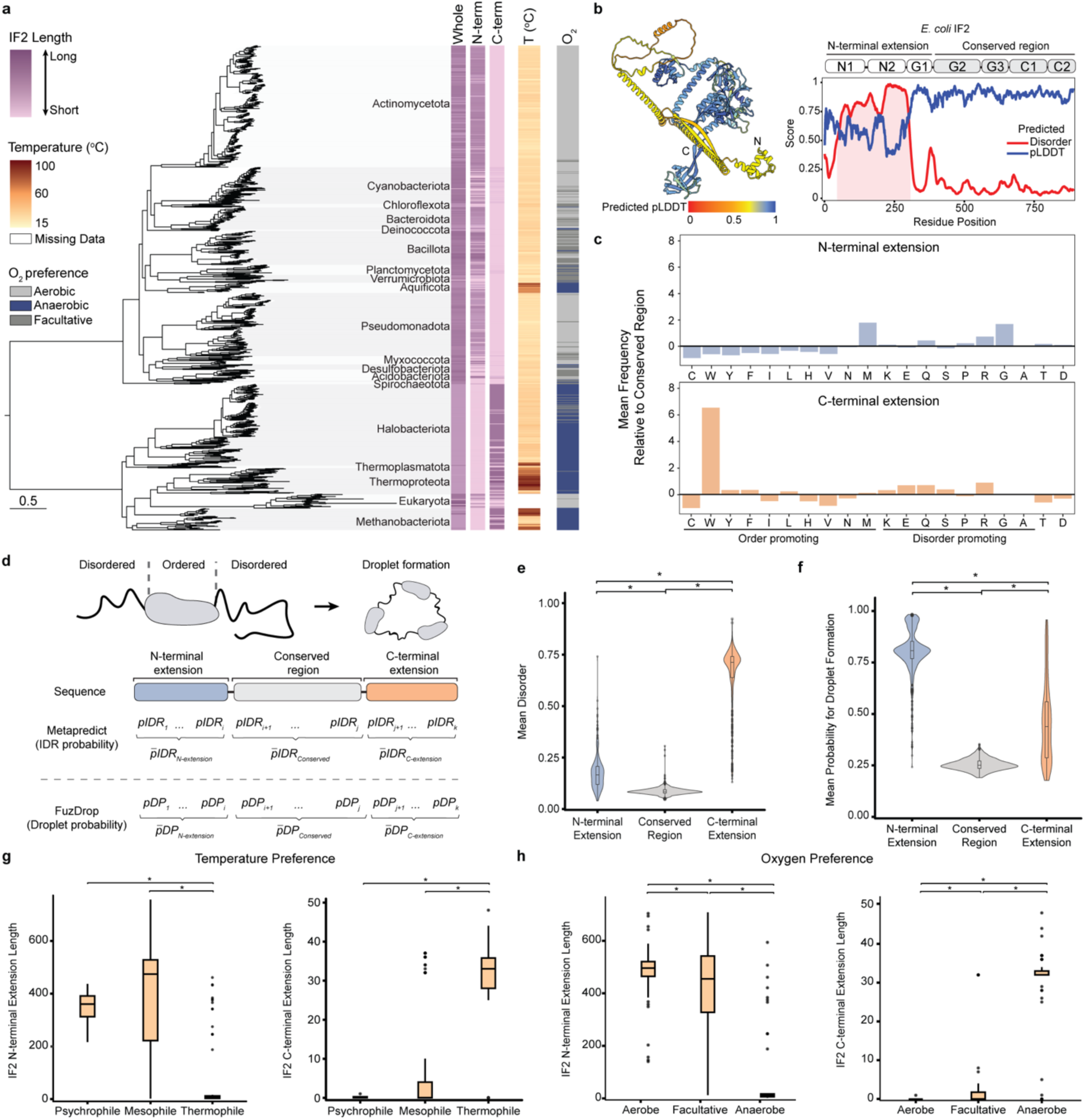
The variation of IF2 terminal regions and the assessment of their intrinsic disorder propensity and correlations to organisms’ environmental preferences. **a)** The species tree with the extension lengths are shown on a color gradient from short (pink) and long (purple) extension lengths. The predicted optimal growth temperature (OGT) of each organism is shown from 15°C (yellow) to 100°C (dark red). Oxygen tolerance is indicated as aerobic (light gray), anaerobic (dark blue), or facultative (dark gray). **b)** Predicted disorder (red) and pLDDT (blue) scores of *E. coli* IF2. The regions with a disorder score higher than 0.3 are shown in red shade. The pLDDT values are projected on the predicted structure of *E. coli* IF2 where low confidence values are shown with red-yellow, high confidence shown in blue. **c)** Mean frequency of amino acids classified as order-promoting and disorder-promoting in IF2 N-terminal extensions (blue) and C-terminal extensions (orange) with respect to the conserved internal region of IF2 sequences. **d)** Representation of disordered and ordered regions on a protein and how they promote phase separation. An overview of calculations for disorder propensities and droplet formation probabilities is shown. **e)** Violin plots comparing mean disorder and **f)** mean probability for droplet (phase-separated condensate) formation between N-terminal extensions (blue), internal conserved regions (gray), and C-terminal extensions (orange). **g)** Boxplots showing the distribution of IF2 N-terminal (left) and C-terminal (right) extension lengths across organisms categorized by temperature preference: psychrophiles, mesophiles, and thermophiles. **h)** Boxplots showing the distribution of IF2 N-terminal (left) and C-terminal (right) extension lengths across organisms with different oxygen preferences: aerobes, facultative, and anaerobes. Asterisks indicate statistically significant differences by Wilcoxon rank-sum test (*p-*value<0.05).

Sequence alignment and domain comparison revealed a conserved catalytic framework consisting of the GTP-binding (G2)^29^, ribosome-interaction (G3)^30^, a linker domain (C1)^31^ and tRNA-binding (C2)^32^ domains, flanked by variable terminal regions (**Fig. 1a, Extended Data Fig. 1**). These core domains display strong sequence conservation across all three domains of life, consistent with previous evidence that the IF2/aIF5B/eIF5B core was already established before the Last Universal Common Ancestor (LUCA)^33^. In contrast, the terminal extensions show extensive lineage-specific diversification (**Fig. 1a**) suggesting plausible additional roles. For example, the N-terminal extension is markedly longer and more variable in bacteria (102–756 amino acids) or eukaryotes (423–979 amino acids) than in archaea (2–21 amino acids). In contrast, C-terminal extensions are short or absent in most bacteria (0–8 aa) but consistently present in a/eIF5B (25–48 amino acids) (**Fig. 1a**). These contrasting length distributions indicate that, after the establishment of the universal IF2 core, terminal extensions arose and diversified independently within major lineages, giving rise to distinct architectural IF2 signatures across the tree of life.

We calculated site-specific selection pressure (dN/dS, also shown as *ω*) across domains using site model-based analysis. Across the full dataset, the mean *ω* for the conserved core (G2–C2 domains) was 0.08 ± 0.1, consistent with strong purifying selection acting to preserve catalytic GTPase, tRNA-binding and ribosome-binding functions (**Extended Data Fig. 1**). In contrast, the N-terminal extensions in bacteria displayed substantially higher *ω* values (0.44 ± 0.4 on average), indicative of neutral evolution, but with localized clusters of purifying sites (*ω* < 0.2) in all bacterial and eukaryotic clades (**Extended Data Fig. 1**). These constrained pockets may correspond to residues involved in secondary structural elements or partner interactions.

For the C-terminal extensions in a/eIF5B, *ω* values are uniformly low (0.07 ± 0.2), with evidence for purifying selection across nearly all sites. This pattern suggests that, once established, these C-terminal regions became integral to the translation initiation complex and have remained under stabilizing constraint. Together, the dN/dS analyses indicate that the IF2 core is under purifying selection, whereas the terminal extensions evolve under mixed regimes of relaxed constraint with lineage-specific conservation in bacteria, and widespread purifying selection in a/eIF5B. These patterns support a model in which the conserved core was established early in the evolution of bacterial initiation and the extensions diversified after the establishment of the ancestral IF2 scaffold, with subsequent selective maintenance of lineage-specific functional elements.

### The disorder and phase separation propensity of IF2 extensions

The N-terminal region of IF2 remains structurally unresolved^12,34–36^. We therefore used structure prediction approach to examine N-terminal extension of *E. coli* IF2. Predictions of full-length *E. coli* IF2 indicate that the N-terminal extension is predominantly unstructured, a characteristic feature of intrinsically disordered protein regions^37^ **(Fig. 1b**). To test whether this is a conserved behavior, we investigated the disorder propensity across all IF2 proteins first by analyzing the amino acid composition of extensions with respect to the conserved internal region. Specifically, we quantified the frequency of amino acids that are known to promote order (C, W, Y, F, I, L, H, V, N, M) and disorder (K, E, Q, S, P, R, G, A)^38^. We showed that extension regions, particularly N-terminal extensions, are more enriched with the disorder-promoting amino acids in comparison to conserved internal region **(Fig. 1c).** Consistently, residue specific disorder predictions confirmed higher intrinsic disorder propensities in both N- and C-terminal extensions relative to the conserved internal region (*p*-value < 2.2 × 10^−16^, Student’s two-sample *t*-test) **(Fig. 1d-e).** Intrinsically disordered regions are often associated with the ability to promote liquid-liquid phase separation (LLPS) within the cell^39–41^. To test whether this behavior applies to IF2 extensions, we quantified the phase separation propensity (i.e., droplet promotion probability) across all IF2s (**Fig. 1d**). Both N- and C-terminal extensions show higher droplet formation probabilities relative to the internal region (*p*-value < 2.2 × 10^−16^, Student’s two-sample *t*-test) **(Fig. 1f)**. Together, these results indicate that the extensions are intrinsically disordered and likely contributed to phase separation.

### IF2 extension length correlations with environmental preferences

Because environmental conditions directly influence protein function and stability, we hypothesized that the diversity in IF2 extensions might be adaptations to environmental conditions of host organism. To test whether IF2 extension diversity reflects adaptations to temperature, we predicted the optimal growth temperature (OGT) of organisms in our dataset using a regression-based model applicable to bacteria and archaea^42^. Our predictions ranged between 16–100°C (**Fig. 1a**), including 5 psychrophiles (OGT ≤ 20°C), 861 mesophiles (20°C < OGT < 45°C), and 107 thermophiles (OGT ≥ 45°C). To assess reliability of predictions, we compared these predictions with curated data from BacDive database^43^, which included 431 organisms with known temperature preferences: 8 psychrophiles, 358 mesophiles, and 65 thermophiles (**Extended Data Fig. 2**). Both predicted and curated datasets revealed that IF2 N-terminal extensions are generally longer in psychrophiles than in thermophiles, with mesophiles exhibiting a broad range of N-terminal sequence lengths (**Fig. 1g, Extended Data Fig. 2**). In contrast, IF2 C-terminal extensions are longer in thermophiles than in psychrophiles or mesophiles (**Fig. 1g, Extended Data Fig. 2**). We showed that organisms’ optimal growth temperature had a weak negative (Pearson *R*=-0.4, *p*-value<0.001; linear regression *R*^2^=0.17, *p*-value<0.001) and a weak positive correlation (Pearson *R*=0.46, *p*-value<0.001; linear regression *R*^2^=0.21, *p*-value<0.001) with N-terminal length and C-terminal length, respectively (**Extended Data Fig. 2**). To assess whether the correlations are influenced by shared evolutionary history, we used Pagel’s lambda (*λ*) phylogenetic regression^44^. Lambda values were found close to 1 (*λ* = 0.99) demonstrate that IF2 extension length is largely constrained by evolutionary history, with no evidence for direct adaptation to host growth temperature (**Extended Data Fig. 2**).

We next examined whether IF2 extensions exhibit evidence of adaptations to oxygen availability in the host environment. We predicted oxygen preferences using Oxyphen which classifies organisms as aerobic/anaerobic/facultative based on the number of O_2_-utilizing enzymes^45^. Organisms having ≥29 O_2_-utilizing enzymes were classified as aerobic, ≤13 as anaerobic, and intermediate cases as facultative^45^. Our dataset included 526 aerobic, 293 anaerobic, and 184 facultative organisms. For validation, we compared predictions with known data from BacDive database covering 153 aerobic, 82 anaerobic, and 26 facultative species. Both predicted and curated data showed that IF2 N-terminal extensions are longer in aerobic and facultative organisms than in anaerobes (**Fig. 1h, Extended Data Fig. 2**). Conversely, C-terminal extensions were consistently longer in anaerobes than in aerobic or facultative organisms across both datasets (**Fig. 1h, Extended Data Fig. 2**). Regression analyses revealed weak positive correlations between N-terminal length and the number of O_2_-utilizing enzymes (Pearson *R*= −0.5, *p*-value<0.001; linear regression *R*^2^=0.25, *p*-value<0.001), and weak negative correlations for C-terminal length (Pearson *R*=0.5, *p*-value<0.001; linear regression *R*^2^=0.25, *p*-value<0.001) (**Extended Data Fig. 2**). As with temperature, phylogenetic regression indicated that these correlations are strongly influenced by shared ancestry (*λ* = 0.97), suggesting that oxygen preference is not a primary driver of IF2 extension length variation. Although IF2 extension length shows broad associations with temperature and oxygen preference across lineages, phylogenetic analyses indicate that these patterns primarily reflect shared evolutionary history rather than direct adaptation to environmental conditions.

### The impact of truncation and swap of IF2 N-terminal extensions on translation and growth

We examined the effect of truncating the N-terminal region of IF2 on translation efficiency. We removed the first 294 amino acids from the N-terminal (NTD Δ1-294 IF2, called ΔN IF2 from here) and compared its activity to full-length IF2 (called WT IF2 from here) (**Fig. 2a**). Both WT IF2 and ΔN IF2 were purified and added at equal concentrations to the *in vitro* transcription-translation system lacking endogenous IF2 (**Fig. 2a, see Materials and Methods**). Translation efficiency was assessed by measuring the activity of luciferase synthesized by the *in vitro* system. Following translation, the luciferase substrate D-luciferin was added, and luminescence was measured as a proxy for luciferase activity, reflecting the efficiency of translation (**Fig. 2a**). We observed that ΔN IF2 supported only half the activity observed with WT IF2 (mean WT IF2=1, mean ΔN IF2= 0.51, Student’s two-sample *t*-test *p*-value=0.003) (**Fig. 2b**). This indicates that the first 294 amino acids are indispensable for translation to occur, but the efficiency significantly decreases to half in *E. coli*.

**Fig 2.**
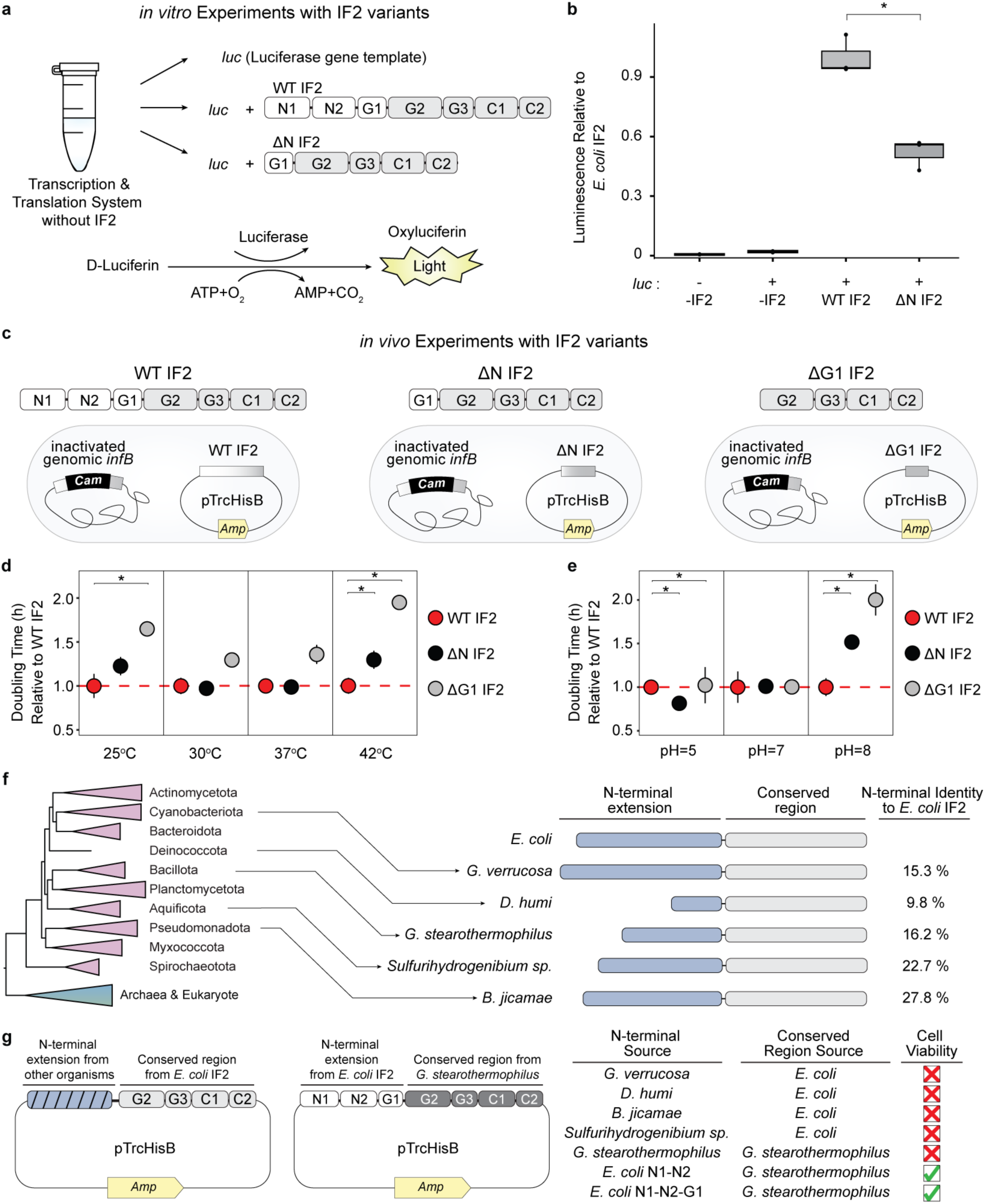
Assessment of the impact of N-terminal truncations of *E. coli* IF2 on translation efficiency and bacterial growth at different conditions**. a)** Overview of in vitro experiments using purified IF2 variants in a customized transcription and translation system without endogenous IF2. Luciferase enzyme synthesized in the in vitro system is used a reporter for translation efficiency. **b)** Luminescence measurements of in vitro reactions relative to the reaction with WT IF2. **c)** The overview of strains and IF2 variants used in the in vivo study**. d)** Doubling time at various temperatures (25 °C, 30 °C, 37 °C, 42 °C) is shown relative to the WT IF2 at that temperature. **e)** Doubling time at various pH (pH 5, pH 7, pH 8) is shown relative to the WT IF2 at that pH. *E. coli* with wild-type IF2 (WT IF2) is shown as red circle; N1-N2 domain truncated IF2 (ΔN IF2) as black circle; and N1-N2-G1 domain-truncated IF2 (ΔG1 IF2) as white circle. Each point represents the distribution of relative doubling time of five replicates (n=5). Asterisks indicate statistically significant differences by Wilcoxon rank-sum test (*p*-value<0.05). The red dashed line indicates WT IF2 doubling time at 1 as reference. **f)** The simplified tree representing bacterial clades. Representative bacterial organisms with IF2 N-terminal extensions were selected and aligned with *E. coli* IF2. The sequence identity of N-terminal extension regions of selected sequences showing the sequence diversity in this region. *G. verrucosa*: *Gloeotece verrucosa*; *D. humi*: *Deinococcus humi*; *G. stearothermophilus*: *Geobacillus stearothermophilus*; *B. jicamae*: *Bradyrhizobium jicamae, Sulfurihydrogenibium sp.: Sulfurihydrogenibium sp.* YO3AOP1. **g)** Overview of construction of strains containing mutant IF2 with N-terminal extensions from different organisms. Additional strains were constructed using the conserved region from *G. stearothermophilus* IF2 and *E. coli* IF2 N-terminal extension. The cell viabilities were indicated as non-viable in red and viable in green.

We next experimentally assessed the impact of IF2 truncation on bacterial growth. Experiments were performed in a specifically constructed *E. coli* SL598R strain, where the chromosomal *infB* gene is disrupted by insertion of chloramphenicol resistance (*CmR*) cassette (**Fig. 2c**) and functional *infB* is supplied via lysogenic λ phage^15^. The λ phage can be selectively excised with a heat pulse at 42°C, making the organism dependent on any IF2 variant provided with plasmid^15^. Using this system, we measured the growth rates of IF2 N-terminal truncation variants under multiple conditions to assess the contribution of the N-terminal region to stress responses. We analyzed three IF2 variants with different N-terminal lengths: full-length *E. coli* IF2 (WT-IF2), an IF2 lacking 1-294 amino acids (ΔN-IF2) and a new variant lacking 1-387 amino acids (NTD Δ1-387 IF2, hereafter ΔG1-IF2) (**Fig. 2c**). For the WT reference, we used *E. coli* MG1655. Growth equivalence between IF2 expressed from the MG1655 genome and from a plasmid in the SL598R background was confirmed (**Extended Data Fig. 3, see Materials and Methods**).

We first assessed the growth of *E. coli* strains expressing WT-IF2, ΔN-IF2, or ΔG1-IF2 across a temperature range from 25°C to 42°C. At 30°C and 37°C, ΔN-IF2 supported growth rates indistinguishable from WT IF2, whereas ΔG1-IF2 reduced growth by ∼25% (**Fig. 2d, Extended Data Fig. 4**). When more extreme temperatures (25°C or 42°C), ΔN-IF2 exhibited a ∼25% reduction in growth relative to WT IF2, while ΔG1-IF2 showed a more pronounced ∼50% decrease (**Fig. 2d, Extended Data Fig. 4**).

We next examined growth under varying pH conditions (pH 5, 7 and 8). Under acidic conditions (pH 5), ΔN-IF2 displayed a 20% reduction in growth relative to WT IF2 (**Fig. 2e, Extended Data Fig. 4**). The effects of N-terminal truncation were more pronounced under basic conditions: at pH 8, ΔN-IF2 reduced growth by ∼30%, and ΔG1-IF2 by ∼50% (**Fig. 2e, Extended Data Fig. 4**).

Together, these results indicate that the G1 domain is required for robust growth across all tested conditions, whereas the N1–N2 regions become particularly important under environmental stress. Thus, in *E. coli*, the IF2 N-terminal region contributes to growth by enhancing tolerance to temperature and pH fluctuations.

To further probe the functional specificity of IF2 N-terminal extensions, we swapped *E. coli* N-terminal region with corresponding segments from diverse bacterial homologs—sharing only 9 to 27% sequence identity (**Fig. 2f**). All such chimeras were nonviable, indicating strong host specificity of IF2 N-terminal domains (**Fig. 2g**). Consistently, complete replacement of *E. coli* IF2 with the homolog from *Geobacillus stearothermophilus* also resulted in loss of viability, whereas chimeric proteins retaining the *E. coli* IF2 N-terminal extension fused to a heterologous conserved core were viable and showed no significant growth defects relative to wild-type IF2 (**Fig. 2g**; **Extended Data Fig. 5**). Together, these results indicate that IF2 N-terminal extensions are functionally specialized to their host background and that introduction of heterologous N-terminal regions is detrimental to cellular viability.

### The impact of C-terminal extensions of IF2 on bacterial growth at different temperature, pH, and under anaerobiosis

Our bioinformatic analysis predicts that several archaeal species in our dataset (particularly the thermophiles) encode C-terminal extensions in IF2 that are generally absent from bacterial homologs (**Fig. 1g-h**). To investigate whether these archaeal-derived C-terminal extensions contribute to stress regulation, especially under varying temperature, pH, and oxygen availability, we selected seven representative candidates (five archaeal, one bacterial, and one fungal) (**Fig. 3a**). We fused each C-terminal extension to the WT IF2 in pTrcHisB plasmid transformed to SL598R strain and evaluated growth under three temperatures (30°C, 37°C, and 42°C), three pH conditions (pH 5, 7, and 8), as well as anaerobic conditions (**Fig. 3b-c, Extended Data Fig. 5**). Addition of the C-terminal extensions enhanced growth at elevated temperature (42°C), with *S. osmophilus* C-terminal extension-containing IF2 exhibiting 23% faster doubling times relative to the WT strain (**Fig. 3c**). Under anaerobic conditions, C-terminal extension variants showed faster doubling time, growing 20-40% faster than the wild type (**Fig. 3c**). In contrast, these variants exhibited impaired growth at lower temperatures (e.g., 30°C), with doubling times 5-30% slower than the wild-type control (**Fig. 3c**). Because the C-terminal extensions were selected from archaeal species that naturally inhabit high-temperature and low-oxygen environments, we speculate that these domains contribute to adaptation to anaerobiosis and thermal stress (**Fig. 3c**). We also observed modestly improved growth of C-terminal extension variants under alkaline conditions, with increases of 0.3 to 11% relative to the WT at pH 8, but not under acidic conditions (**Extended Data Fig. 6**). Together, these findings suggest that archaeal IF2 C-terminal extensions can enhance bacterial fitness under specific environmental stresses and may function as conserved stress-responsive modules.

**Fig 3.**
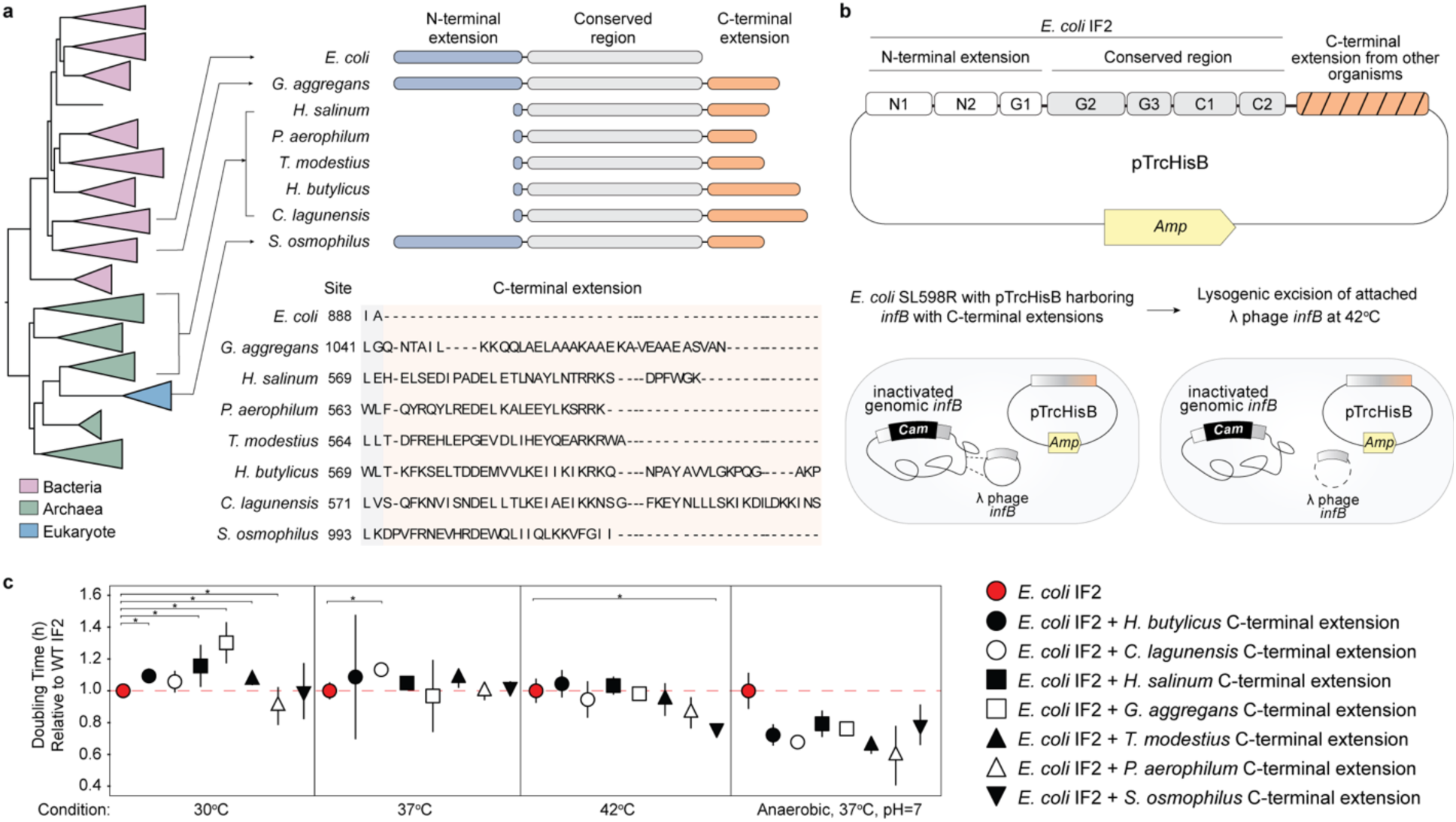
The impact of addition of C-terminal extensions to *E. coli* IF2 on bacterial growth. **a)** The simplified tree representing bacterial, archaeal and eukaryotic clades in pink, green and blue, respectively. Representative organisms with IF2 C-terminal extensions were selected and aligned with *E. coli* IF2. The alignment zooms in C-terminal extension regions of selected sequences showing the sequence diversity in this region. **b)** Overview of construction of strains containing mutant IF2 with C-terminal extensions from different organisms. **c)** Doubling time at various temperatures (30°C, 37°C, 42°C) and under anaerobic condition are shown relative to the WT IF2. Each point represents the distribution of relative doubling time of five replicates (n=5), except the data for anaerobic condition (n=3). Asterisks indicate statistically significant differences by Wilcoxon rank-sum test (*p*-value<0.05). The red dashed line indicates WT IF2 doubling time at 1 as reference. *H. butylicus*: *Hyperthermus butylicus*; *C. lagunensis*: *Caldisphaera lagunensis*; *H. salinum*: *Halorarum salinum*; *G. aggregans*: *Granulicella aggregans*; *T. modestius*: *Thermocladium modestius*; *P. aerophilum*: *Pyrobaculum aerophilum*; *S. osmophilus*: *Schizosaccharomyces osmophilus*.

### Structural diversity of IF2 extensions across three domains of life

Our experimental results indicate that IF2 terminal extensions exhibit distinct functional behaviors, yet their poor sequence conservation makes it difficult to infer the molecular basis of these differences from sequence analysis alone. In contrast, recurrent secondary-structure architectures that diversify across lineages may provide greater explanatory power^46^. To identify structural features within IF2 extension regions, we predicted the structures of 809 IF2 homologs, followed by clustering using Foldseek^47^ and manual curation.

This analysis identified seven distinct IF2 structural types, defined by the organization and secondary-structure composition of α-helical and helix–turn–helix (HTH) elements within the terminal regions (**Table 1**, **Fig. 4**). The distribution of these architectures is strongly lineage biased. Bacterial IF2s encompass five structural types (Types 2–6), with Type 4, characterized by a double-HTH motif followed by a long coil, being the most prevalent. Archaeal IF5B is represented exclusively by Type 1, which contains short C-terminal α-helices, whereas eIF5B corresponds to Type 7, featuring extended α-helical termini (**Table 1**, **Fig. 4, Extended Data Fig. 7**).

**Fig 4.**
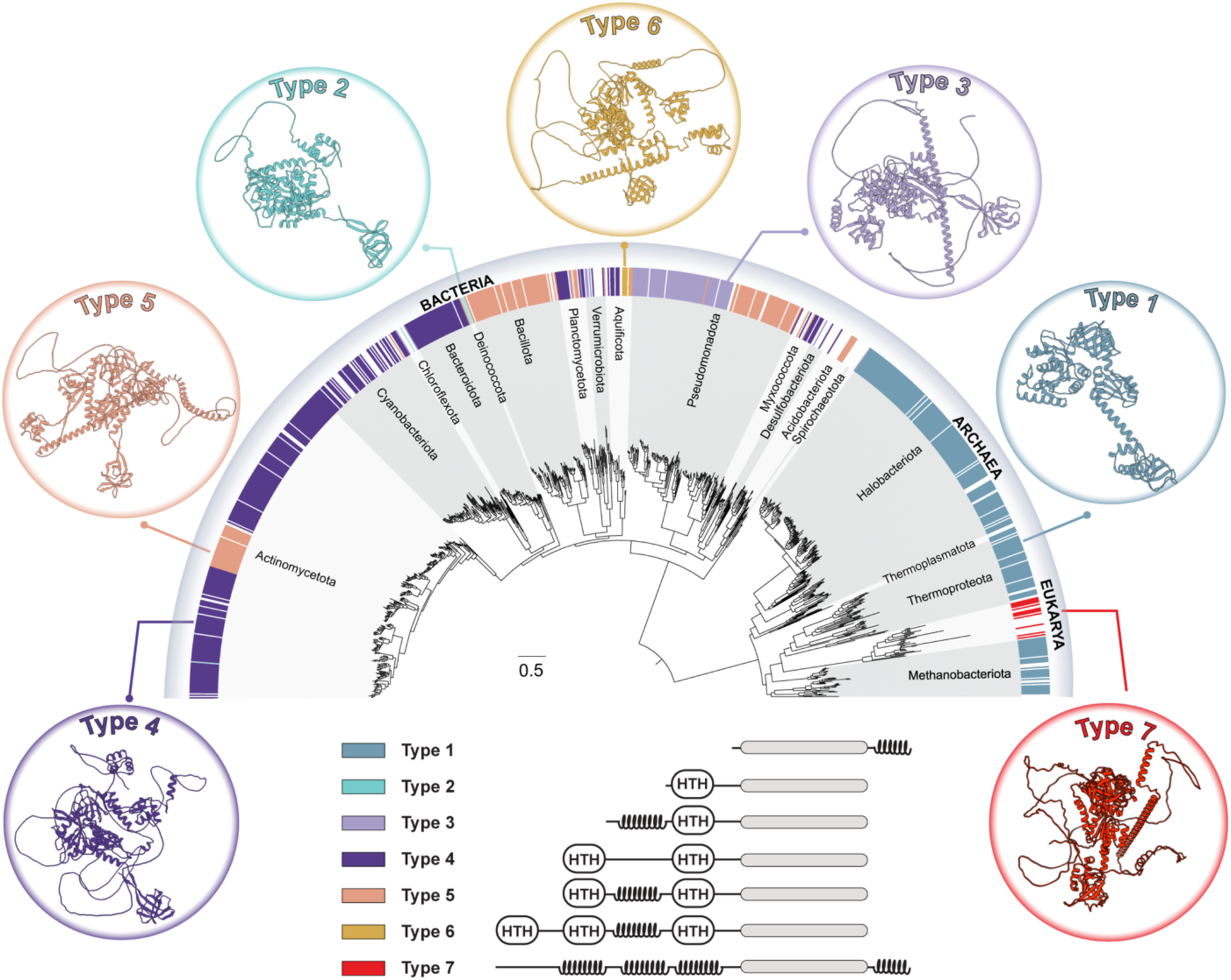
Structural diversity of IF2 extensions across three domains of life. A species tree was constructed with ten ribosomal proteins from 1003 representative organisms. The tree is shaded based on the taxonomic clades. Seven IF2 types were determined based on the structures of N-terminal and C-terminal extensions, and one representative predicted structure is shown for each IF2 type in circles. HTH: Helix-turn-helix.

**Table 1.** The list of defined IF2 types with key structural features, taxonomic distribution, relative frequency in the dataset and representative example.

| IF2 Type | Key Structural Features | Taxonomic Distribution | Freq (%) | Representative Example |
| --- | --- | --- | --- | --- |
| Type 1 | Conserved G2–C2 catalytic core with additional short α-helices appended at the C-terminus | Halobacteriota<br>Methanoproteota<br>Thermoplasmatota<br>Thermoproteota | 28 | *Methanosarcina acetivorans* |
| Type 2 | N-terminal with one Helix-Turn-Helix (HTH) motif preceding the conserved G2-C2 catalytic core | Deinococcota | 0.6 | *Deinococcus humi* |
| Type 3 | N-terminal with long α-helices and one Helix-Turn-Helix (HTH) motif preceding the conserved G2-C2 catalytic core | Pseudomonadota<br>Chloroflexota<br>Planctomycetota<br>Verrucomicrobiota | 10 | *Mesorhizobium japonicum* |
| Type 4 | N-terminal with two Helix-Turn-Helix (HTH) motifs separated by a long unstructured coil preceding the conserved G2-C2 catalytic core | Actinomycetota<br>Bacillota<br>Cyanobacteriota | 39 | *Streptomyces qaidamensis* |
| Type 5 | N-terminal with two Helix-Turn-Helix (HTH) motifs separated by long α-helices preceding the conserved G2-C2 catalytic core | Actinomycetota<br>Bacteroidota<br>Pseudomonadota | 19 | *Paraburkholderia edwinii* |
| Type 6 | N-terminal with three Helix-Turn-Helix (HTH) motifs separated by α-helical linkers preceding the conserved G2-C2 catalytic core | Aquificota | 0.6 | *Sulfurihydrogenibium azorense* |
| Type 7 | Extended α-helical regions at both N- and C-termini in addition to the G2-C2 catalytic core | Ameobozoa<br>Fungi<br>Metazoa<br>Sar<br>Viridiplantae | 1.5 | *Saccharomyces pombe* |

Given the extensive structural diversity of bacterial IF2, we classified the homologs used in the N-terminal swap experiments (**Fig. 2f-g**). These proteins span multiple IF2 structural types: *D. humi* (Type 2), *B. jicamae* (Type 3), *G. verrucosa* (Type 4), *E. coli* and *G. stearothermophilus* (Type 5) and *Sulfurihydrogenibium sp.* (Type 6). Together, with the results of the N-terminal swap experiments (**Fig. 2f-g**), this classification demonstrates that IF2 N-terminal extensions are not functionally interchangeable across distinct structural IF2 types.

### The correlation test between bacterial IF2 N-terminal extension and NusA interactions

A recent study demonstrated that the transcription elongation factor NusA interacts transiently and isoform-specifically with IF2 through the IF2α N-terminal Domain I and the KH2/AR1 domains of NusA^48^. NusA is a conserved bacterial transcription factor that regulates transcription termination by modulating RNA polymerase pausing and stabilizing terminator RNA structures through its modular domain architecture^49–51^. Its C-terminal region, comprising an S1 domain followed by tandem KH and acidic repeat (AR) motifs, mediates interactions with nascent RNA and regulatory proteins^49–51^. Notably, the interaction between IF2 N-terminal extension and the KH2/AR1 domains of NusA occurs frequently *in vivo* and is enriched near the nucleoid, without altering the core mechanism of translation initiation. Instead, NusA is proposed to spatially concentrate IF2 near nascent transcripts, thereby facilitating transcription–translation coupling. Given this functional association, we tested whether the length of the IF2 N-terminal extension lhas co-evolved with variation in the C-terminal region of NusA.

Across bacterial species in our IF2 dataset, we identified corresponding NusA homologs and assessed the relationship between IF2 N-terminal extension length and NusA C-terminal extension length. This analysis revealed a weak but statistically significant negative correlation between the two features (Pearson’s r = −0.18, *p* = 1.5 × 10⁻⁶) (**Extended Data Fig. 8**).

## Discussion

We examined the evolutionary diversity and functional significance of IF2 terminal extensions by integrating comparative genomics, targeted functional experiments, structural prediction and environmental correlations. Our findings reveal that IF2 extensions represent a previously underappreciated layer of adaptive diversification, superimposed on a highly conserved translational GTPase, contributing to lineage-specific responses to environmental conditions across the tree of life.

Comparative analyses show substantial variation in IF2 terminal length, particularly within bacterial N-terminal extensions. Consistent with previous studies, these the N-terminal regions are dispensable for growth under optimal laboratory conditions^15–17,29,52^. However, the IF2 N-terminus has also been implicated in chaperone activity, DNA binding, and cold-shock response^52–55^, suggesting regulatory roles beyond canonical initiation. The absence of homologs extensions in archaeal IF5B further supposed the idea that bacterial and archaeal IF2 extensions arose independently following divergence from an ancestral IF2 lacking terminal additions^33^. Together, these observations indicate that IF2 extensions did not evolve to preserve a universal function but instead emerged as lineage-specific solutions layered onto a conserved catalytic core (**Fig. 5**).

**Fig 5.**
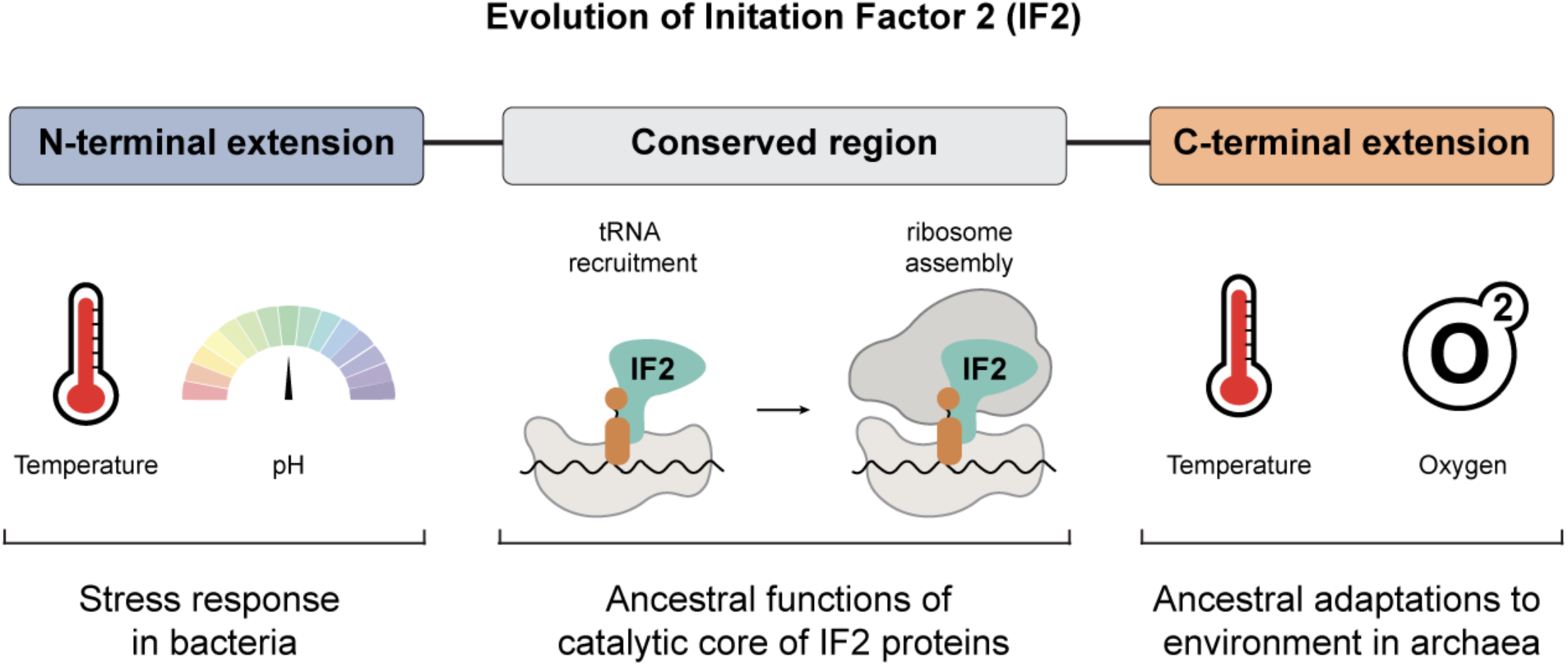
Schematic representation of the proposed role of IF2 in bacterial stress adaptation. The conserved core of IF2 functions in translation initiation complex assembly, whereas the N-terminal region and C-terminal extensions contribute to adaptation to elevated temperature and alkaline pH. In addition, the C-terminal extensions are required for adaptation to anaerobiosis.

Structural predictions reveal that IF2 extensions are predominantly intrinsically disordered, consistent with their absence from high-resolution structures. Intrinsically disordered regions are known to facilitate transient interactions, confer conformational flexibility and enable responsiveness to environmental change, including through liquid–liquid phase separation (LLPS)^39–41,56–58^.

Across all IF2 extension types, we observed enrichment of disorder-promoting residues and predicted phase-separation potential, suggesting that these features are intrinsic to extension regions rather than confined to specific lineages (**Fig. 1b-f**). Although these predictors are consistent with regulatory role for IF2 extensions, direct observation of their biophysical behavior *in vivo* will be required to establish their mechanistic contributions.

While variation in translation initiation mechanisms have been examined in the context of growth conditions^59^, individual translation factors have not been systematically analyzed from an environmental adaptation perspective. For example, environmental tuning has been reported for specific ribosomal proteins^60^, but IF2 has not previously been considered in this framework. Our analyses indicate that IF2 extension length co-varies with broad lifestyle traits (**Fig. 1g-h**) but not with specific contemporary ecological parameters (**Extended Data Fig. 2**). This pattern suggests that IF2 extensions may reflect ancient selective pressures associated with major lineage divergence, rather than fine-scale optimization to present-day niches (**Fig. 5**).

Functional assays show that deletion of the IF2 N-terminal extension reduces translation efficiency *in vitro* but causes only modest growth defects *in vivo* under optimal conditions, consistent with prior observations^12,16,17^. (**Fig. 2b, 2d-e**). The discrepancy suggests that compensatory interactions within the cellular environment partially buffer the loss of the N-terminal region. Under environmental stress, however, including non-optimal temperatures and altered pH— loss of the IF2 N-terminus resulted in pronounced growth defects (**Fig. 2d-e**). These findings extend earlier links between the IF2 N-terminus and cold-shock response^52,61^ and support a broader role in stress adaptation. In addition, the strong host specificity observed in N-terminal swap experiments indicate that these regions may co-evolve with interacting partners, contributing to species-specific translation regulation^62–64^. Although we tested co-variation between IF2 N-terminal length and the C-terminal region of NusA, the weak correlation observed suggests that additional interaction networks remain to be explored.

In contrast to bacterial IF2, little is known about the function of C-terminal extensions present in archaeal and eukaryotic homologs. By introducing these extensions into a bacterial context, we found that archaeal-derived C-terminal regions enhanced *E. coli* growth under conditions resembling archaeal native environments, including elevated temperature, alkaline pH, and anaerobiosis (**Fig. 3b, c; Extended Data Fig. 6**). These benefits were accompanied by reduced fitness under mesophilic conditions, consistent with trade-offs associated with environmental specialization. Together, these results support a model in which archaeal IF2 C-terminal extensions function as modular elements that tune translation initiation under extreme conditions, contributing to ancestral niche specialization in archaea (**Fig. 5**). Their absence from most bacterial IF2 homologs likely reflects divergent selective pressures on extensions rather than altering the functional compatibility of the conserved initiation core.

Protein synthesis is among the most energy-intensive cellular processes, requiring precise coordination to balance the efficient maintenance of cellular currency with functional responsiveness^65,66^. Our findings suggest that IF2 terminal extensions contribute to this balance by modulating translation under stress without altering the conserved catalytic mechanism. By further classifying IF2 into distinct structural types based on extension architecture, this study provides a framework for understanding how translation factors can diversify to meet environmental demands while preserving core functionality.

## Materials and Methods

### Data collection and phylogeny construction

We filtered genomes from the GTDB (v2.4.0)^67^ by selecting those with >95% completeness and <5% contamination, resulting in approximately 25,000 bacterial and 2,700 archaeal genomes. For eukaryotes, we included 121 complete genomes obtained from NCBI RefSeq Genomes. After downloading the corresponding proteome files for each genome using NCBI *eutils*^68^, we constructed a local protein database with *makeblastdb*. To build a species tree, we searched for ten universally conserved ribosomal proteins (uS3, uS8, uS17, uS19, uL2, uL3, uL5, uL6, uL14, uL16) within the local database using *blastp* with the following parameters*: -evalue 1e-5 - word_size 5 -matrix BLOSUM62 -max_target_seqs 100000*. We removed duplicate entries and non-target sequences from the BLAST hits, retaining a total of 1003 genomes including 701 bacteria, 272 archaea, and 30 eukaryotes. Each ribosomal protein was aligned individually using MAFFT L-INS-I (v7.90)^69^, and the resulting alignments were concatenated. We inferred the species tree using IQ-TREE (v1)^70^ with the LG+R10 substitution model selected by ModelFinder^71^ and rooted the tree using the minimal ancestor deviation (MAD) method^72^.

### IF2 sequence and structure analysis

We searched for IF2/IF5B homologs in the 1003 genomes used for species tree construction using *blastp* with the same parameters as above. We excluded hits annotated as hypothetical or putative, as well as mitochondrial IF2 sequences from eukaryotes. This filtering resulted in a final set of 809 IF2 sequences, comprising 563 bacterial IF2, 234 aIF5B, and 12 eIF5B. The IF2 homologs were aligned using MAFFT L-INS-I (v7.90)^69^. Using the *E. coli* IF2 domain boundaries as a reference^29^, we annotated each IF2 domain in the alignment. The regions upstream of the GTPase domain (starting at residue 388) were defined as N-terminal extensions, while regions downstream of the C-terminal domain (ending at residue 890) were considered C-terminal extensions. To assess sequence length variability, we normalized the length of each domain in individual sequences by dividing it by the corresponding domain length in the full IF2 alignment. We predicted the structures of all 809 IF2 proteins using ColabFold^73^ and clustered the predicted structures using FoldSeek^47^. After manual inspection of the resulting clusters, we identified seven distinct structural types of IF2. The list of structural types of each IF2 homologs are shown in **Extended Data 1**.

### Selection analysis

We collected the nucleotide sequences corresponding to the IF2 proteins and aligned them to the reference protein alignments to generate codon alignments using PAL2NAL^74^. To optimize computational efficiency, we created separate codon alignments for taxonomic groups containing at least 10 organisms; for smaller groups, we combined closely related sister clades. We then estimated the ratio of non-synonymous to synonymous substitutions (dN/dS or *ω*) using different site models in PAML v4.9^75^: *i*) M0, which assumes a single ω across all sites (null model); *ii*) M1, the nearly neutral model (*ω₀*<1; *ω₁*=1); and *iii*) M2, the positive selection model (*ω₀*<1; *ω₁*=1; *ω₂>*1). We used likelihood ratio tests (LRTs) to determine which model best fit the data.

### Prediction and curation of trait data

Optimal growth temperature (OGT) and oxygen preference data were collected from BacDive^43^ by matching RefSeq accession numbers. Due to the limited availability of experimentally determined trait data, we also predicted OGT and oxygen preference for all genomes. OGT was predicted for both bacterial and archaeal genomes using the OGT_prediction tool^42^. The regression model applied included both Bacteria and Archaea and excluded features based on 16S rRNA gene content and genome size. Based on the predicted temperatures, organisms were classified as psychrophilic (<20°C), mesophilic (20–45°C), or thermophilic (>45°C). Oxygen metabolism was predicted using Oxyphen^45^, which infers oxygen preference based on the abundance of genes encoding oxygen-utilizing enzymes in each genome. Following the thresholds defined in Oxyphen tool^45^, organisms having oxygen utilizing enzymes fewer than 13 were classified as anaerobic, those with more than 29 oxygen utilizing enzymes were classified as aerobic, and those having oxygen utilizing enzymes between 13–29 were classified as facultative. To assess statistical significance (*p*-value < 0.05), we used the Wilcoxon rank-sum test with Benjamini-Hochberg correction implemented in R. The predicted trait data are shared in **Extended Data 2**.

### Correlation analysis

We assessed the relationship between IF2 N-terminal and C-terminal extension lengths and traits (OGT and oxygen preference) using Pearson correlation and linear regression using *cor.test* and *lm* function in R, respectively. To account for biases caused by shared evolutionary history, we performed phylogenetic regression using the *phylolm* function with Pagel’s lambda model from the *phylolm* R package (version 2.6.2)^44^. In this model, the lambda (*λ*) parameter reflects the strength of the phylogenetic signal, ranging from 0 (phylogenetic independence) to 1 (strong phylogenetic dependence).

### Prediction of intrinsic disorder and droplet formation propensity

We investigated disorder propensity by analyzing the amino acid composition of the N-terminal extensions, C-terminal extensions, and the conserved internal region between them. To quantify the composition, we calculated the frequency of amino acids known to promote structural order (C, W, Y, F, I, L, H, V, N, M) and disorder (K, E, Q, S, P, R, G, A)^38^ by dividing the count of each residue by the total length of the corresponding region. Disorder propensity for each region (N-terminal extension, internal region, and C-terminal extension) was predicted using Metapredict^76^, and the mean disorder score was calculated for each region by averaging the per-residue scores. To estimate the probability of phase separation via droplet formation, we used FuzDrop and FuzDrop^77^ with a custom Python script that submitted each sequence to the FuzDrop webserver. We then calculated the mean droplet formation probability for each region by averaging the scores for the residues within the regions.

### Plasmid construction

For protein expression, pET21a(+) plasmids were used as vector backbone. Full length *E. coli infB* gene and *ΔN IF2 (Δ1-294)* genes were cloned into pET21a(+) using HindIII-HF (NEB-R3104S) and EcoRI (NEB-R3101S) restriction enzymes. For *in vivo* experiments, pTrcHisB were used as backbone. *E. coli infB* gene, *ΔN IF2 (Δ1-294)* and *ΔG1 IF2 (Δ1-387)* were inserted into backbone. To construct pTrcHisB-WT IF2, the N1-N2-G1 gene fragment was ordered from Twist Biosciences to clone into pTrcHisB - ΔG1 IF2 using NcoI Fast Digestion enzyme (Thermofisher - FD0574). The N-terminal extensions derived from bacterial species — *Gloeotece verrucosa*, *Deinococcus humi*, *Bradyrhizobium jicamae, Sulfurihydrogenibium sp.* YO3AOP1 — were ordered from Twist Biosciences. The N-terminal extensions were cloned into the pTrcHisB-ΔG1 IF2 by adding the N-terminal sequences right before the ΔG1 IF2 nucleotide sequence. For IF2 derived from *G. stearothermophilus*, the full-length sequence was ordered from Twist Bioscience. The full-length sequence, the ΔN sequence, and the ΔG sequence were used as insert sequences, and the corresponding regions of pTrcHisB were used as backbone sequences, and each was amplified by PCR. Then plasmids were then constructed using Gibson assembly mix (NEB) with these DNA fragments respectively.

The C-terminal extensions were amplified from a synthetic DNA (Twist Bioscences) that was designed in such a fashion that it contained the 7 C-terminal extensions in tandem repeat. Of the seven C-terminal extensions, five were derived from archaeal species — *Hyperthermus butylicus* DSM5456, *Caldisphaera lagunensis* DSM15908, *Halobaculum salinum*, *Thermocladium modestius*, and *Pyrobaculum aerophilum*. One extension originated from a bacterial species (*Granulicella aggregans*), and one from a representative fungus (*Schizosaccharomyces osmophilus*) (**Extended Data 1 and 2**). The C-terminal extensions were added to the pTrcHisB-WT IF2. All cloning were done with Gibson assembly mix (NEB) using the manufacturer instructions. The constructed plasmids were transformed into *E. coli* DH5α chemically competent cells. The plasmids were miniprepped following QIAprep Spin Miniprep Kit protocol. The plasmid constructs were assessed with plasmid sequencing. As with the N-terminal extensions, pTrcHisB-WT IF2 plasmid harboring the C-terminal extensions from 7 representative species listed above were transformed in *E. coli* SL598R strain and lambda-*infB* was excised from the genome using heat curation^15^.

The list of primers and nucleotide sequences that were used in plasmid construction are shared in **Extended Data 3**.

### Protein expression and purification

pET21a(+)-ΔN IF2 and pET21a(+)-WT IF2 were expressed in *E. coli* BL21(DE3). Cultures were grown in LB medium supplemented with carbenicillin (100 µg/mL) to mid-log phase (OD_600_ = 0.3–0.6), then induced with 1 mM IPTG and incubated overnight at 16°C, 250 rpm. Cells were harvested by centrifugation (4500 rpm, 15 min, 4°C) and stored at –80°C until use. For purification, pellets were resuspended in TMN lysis buffer [100 mM Tris HCl pH 7.5, 25 mM MgCl_2_, 1M NaCl, 25% Glycerol, water] and lysed by sonication (30s on, 10s off cycles for ∼30 min). Lysates were clarified by centrifugation at 4000 xg at 4°C for 10 min. The supernatant was subjected to Ni-NTA affinity purification at 4°C. Bound proteins were washed, eluted in 500 µL fractions, and analyzed by SDS-PAGE. Elution were pooled and precipitated overnight using ammonium sulphate. Then further purified by size-exclusion chromatography on a HiLoad™ 26/600 Superdex™ 200 pg column (Cytiva) equilibrated in 1X TMNDN120 buffer [400 mM Tris HCl pH 7.5, 400 mM NH_4_Cl, 800 mM NaCl, 50 mM MgCl_2,_ water]. The eluted fractions were collected and concentrated using 3K Amicon Ultra filter. Protein concentration was determined using the BCA assay (Pierce), with BSA as the standard. The final preparation yielded Purified protein was supplemented with 25% glycerol and 1 mM DTT, aliquoted, and stored at −80°C.

### Luciferase activity assay

For luminescence measurements, customized PURExpress *In Vitro* Protein Synthesis Kit (NEB) lacking endogenous IF2 was used (PURExpress-ΔIF2). The concentrations of purified ΔN IF2 and WT IF2 were normalized on SDS-PAGE for equal concentration of purified proteins (**Extended Data Fig. 9**). Four reactions were prepared: *i*) no DNA template, PURExpress-ΔIF2, no IF2; *ii*) luciferase DNA template (pET21a(+)-*luc*), PURExpress-ΔIF2, no IF2; *iii*) luciferase DNA template (pET21a(+)-*luc*), PURExpress-ΔIF2, WT IF2; *iv*) luciferase DNA template (pET21a(+)-*luc*), PURExpress-ΔIF2, ΔN IF2. All reactions were incubated at 37°C for 4 hours. Three replicates of each reaction were transferred to 96-well white-walled, flat bottom plates. 100 μL of luciferase substrate from Promega Luciferase System (E1500) was added to each reaction. The luminescence was measured at 580 nm for 10 minutes using CLARIOstar Plus Plate Reader while using CLARIOstar Plus pump to confirm that all luciferase reactions were added and measured per the instrument to remove any human error in luciferase assay measurements.

### Constructing mutant strains

The plasmid constructs for N-terminal truncation pTrcHisB-WT IF2, pTrcHisB-ΔN IF2 (Δ1-294), pTrcHisB-ΔG1 IF2 (Δ1-387), N-terminal extension swap (derived from *Gloeotece verrucosa*, *Deinococcus humi*, *Geobacillus stearothermophilus*, *Bradyrhizobium jicamae, Sulfurihydrogenibium sp.* YO3AOP1) and pTrcHisB-WT IF2 harboring 7 C-terminal extensions (derived from *Hyperthermus butylicus* DSM5456, *Caldisphaera lagunensis* DSM15908, *Halobaculum salinum*, *Thermocladium modestius*, *Pyrobaculum aerophilum*, *Granulicella aggregans* and *Schizosaccharomyces osmophilus*) (**Extended Data 3**) were individually transformed to *E. coli* SL598R strains by electroporation at 1700 V for 6 ms. The transformed cells were plated on selection media with chloramphenicol (Cam) and carbenicillin (Carb) antibiotics (LB-Cam-Carb) and incubated at 30°C overnight. The colonies picked from single colony transformants and cultured in LB-Cam-Carb selection media at 30°C overnight. The cultures were diluted to OD_600_ 0.01 in 10 mL and incubated until OD_600_ 0.25-0.3. 1 mM IPTG was added, and the cultures were incubated for 2 hours. After 2 hours, the cultures were diluted to OD_600_ 0.3 in 1.5 mL LB. The cultures were heat-pulsed at 42°C for 5, 10 and 30 minutes. The cultures were chilled on ice and spinned down at 5000 xg for 5 min at 4°C. The pellet was resuspended in 100 μL LB-Carb-Cam and plate on the selection media, incubated at 42°C overnight. For truncated IF2 variants, the excision of genomic *infB* gene was assessed with PCR targeting the N-terminal region. For WT IF2, the excision of genomic *infB* gene was assessed by whole genome sequencing.

### Growth assays

The strains with different IF2 variants were inoculated in LB-Cam-Carb selection media and grown at 30°C overnight. The cultures were diluted to OD_600_ 0.01. Five replicates of 100 μL of each culture were transferred to 96-well flat-bottom plates. The plates were covered with semi-permeable membrane. The growths were measured at different temperatures (25°C, 30°C, 37°C, 42°C) for 24 hours measuring absorbance at every 15 minutes using CLARIOstar Plus Plate Reader. The growth assay was repeated in the same way using LB media with different pH (pH 5, pH 7, pH 8). For growth assays conducted with WT-IF2 harboring 7 different C-terminal extensions were conducted similarly as described above for 3 different temperatures (25°C, 30°C, 37°C, 42°C) and three different pH (pH 5, pH 7, pH 8) except the growth measurements were taken every 20 min. Anaerobic growth assays were conducted in Hungate tubes containing 5 ml of LB-Cam-Carb at 37°C. The data were collected from three biological replicates. All the growth measurements were analyzed in R.

### NusA sequence analysis

Corresponding NusA proteins were identified for each bacterial organism containing IF2 in the dataset using rentrez (v1.2.4)^78^. NusA sequences were aligned with MAFFT L-INS-I (v7.90)^69^. C-terminal extensions were defined as the regions following the last conserved NusA domain, and their lengths were extracted. NusA C-terminal extension lengths were then correlated to IF2 N-terminal extension lengths using Pearson correlation analysis. The accession IDs of NusA proteins used in the analysis are shared in **Extended Data 2**.

## Data availability

All data and R scripts are available on GitHub (https://github.com/evrimfer/if2_extension).

## Acknowledgments

We thank Thomas Dever (NIH) for providing SL598R *E. coli* strains. We thank David Baum, Daniel Amador-Noguez, Michael Thomas and Robert Landick, as well as the participants of the Telluride Workshop on Plasticity in Biological Organization for critical feedback. This work was supported by the John Templeton Foundation (#61926), UW-Madison Bacteriology Department Peterson Predoctoral Fellowship (EF), UW-Madison Alumni Foundation Hiroshi-Sugiyama Fund (EF), and the National Institutes of Health, USA (NIH #T32GM136536) (KM).

## References

1 McCarthy, J. E. G. & Gualerzi, C. Translational control of prokaryotic gene expression. Trends in Genetics 6, 78–85 (1990).

2 Rodnina, M. V. Translation in Prokaryotes. Cold Spring Harb Perspect Biol 10 (2018).

3 Schmitt, E. et al. Recent Advances in Archaeal Translation Initiation. Front Microbiol 11, 584152 (2020).

4 Brito Querido, J., Díaz-López, I. & Ramakrishnan, V. The molecular basis of translation initiation and its regulation in eukaryotes. Nat Rev Mol Cell Biol 25, 168–186 (2024).

5 Pestova, T. V. et al. The joining of ribosomal subunits in eukaryotes requires eIF5B. Nature 403, 332–335 (2000).

6 Kaledhonkar, S. et al. Late steps in bacterial translation initiation visualized using time-resolved cryo-EM. Nature 570, 400–404 (2019).

7 Kazan, R. et al. Role of aIF5B in archaeal translation initiation. Nucleic Acids Res 50, 6532–6548 (2022).

8 Dever, T. E. & Green, R. The elongation, termination, and recycling phases of translation in eukaryotes. Cold Spring Harb Perspect Biol 4, a013706 (2012).

9 Krzyzosiak, A., Pitera, A. P. & Bertolotti, A. An Overview of Methods for Detecting eIF2α Phosphorylation and the Integrated Stress Response. Methods Mol Biol 2428, 3–18 (2022).

10 Marintchev, A. & Wagner, G. Translation initiation: structures, mechanisms and evolution. Q Rev Biophys 37, 197–284 (2004).

11 Milon, P. et al. The ribosome-bound initiation factor 2 recruits initiator tRNA to the 30S initiation complex. EMBO Rep 11, 312–316 (2010).

12 Simonetti, A. et al. Involvement of protein IF2 N domain in ribosomal subunit joining revealed from architecture and function of the full-length initiation factor. Proc Natl Acad Sci U S A 110, 15656–15661 (2013).

13 Atkinson, G. C. The evolutionary and functional diversity of classical and lesser-known cytoplasmic and organellar translational GTPases across the tree of life. BMC Genomics 16, 78 (2015).

14 Fer, E., Yao, T., McGrath, K. M., Goldman, A. D. & Kaçar, B. The origins and evolution of translation factors. Trends Genet 41, 590–600 (2025).

15 Laalami, S., Putzer, H., Plumbridge, J. A. & Grunberg-Manago, M. A severely truncated form of translational initiation factor 2 supports growth of *Escherichia coli*. J Mol Biol 220, 335–349 (1991).

16 Moreno, J. M., Drskjøtersen, L., Kristensen, J. E., Mortensen, K. K. & Sperling-Petersen, H. U. Characterization of the domains of *E. coli* initiation factor IF2 responsible for recognition of the ribosome. FEBS Lett 455, 130–134 (1999).

17 Caserta, E. et al. Ribosomal interaction of *Bacillus stearothermophilus* translation initiation factor IF2: characterization of the active sites. J Mol Biol 396, 118–129 (2010).

18 Choi, S. K. et al. Physical and functional interaction between the eukaryotic orthologs of prokaryotic translation initiation factors IF1 and IF2. Mol Cell Biol 20, 7183–7191 (2000).

19 Marintchev, A., Kolupaeva, V. G., Pestova, T. V. & Wagner, G. Mapping the binding interface between human eukaryotic initiation factors 1A and 5B: A new interaction between old partners. Proceedings of the National Academy of Sciences 100, 1535–1540 (2003).

20 Acker, M. G., Shin, B.-S., Dever, T. E. & Lorsch, J. R. Interaction between Eukaryotic Initiation Factors 1A and 5B Is Required for Efficient Ribosomal Subunit Joining. Journal of Biological Chemistry 281, 8469–8475 (2006).

21 Fringer, J. M., Acker, M. G., Fekete, C. A., Lorsch, J. R. & Dever, T. E. Coupled Release of Eukaryotic Translation Initiation Factors 5B and 1A from 80S Ribosomes following Subunit Joining. Molecular and Cellular Biology 27, 2384–2397 (2007).

22 Zheng, A. et al. X-ray structures of eIF5B and the eIF5B-eIF1A complex: the conformational flexibility of eIF5B is restricted on the ribosome by interaction with eIF1A. Acta Crystallogr D Biol Crystallogr 70, 3090–3098 (2014).

23 Nag, N. et al. eIF1A/eIF5B interaction network and its functions in translation initiation complex assembly and remodeling. Nucleic Acids Research 44, 7441–7456 (2016).

24 Fernández-Pevida, A. et al. The eukaryote-specific N-terminal extension of ribosomal protein S31 contributes to the assembly and function of 40S ribosomal subunits. Nucleic Acids Research 44, 7777–7791 (2016).

25 Derbikova, K. et al. Biological and Evolutionary Significance of Terminal Extensions of Mitochondrial Translation Initiation Factor 3. International Journal of Molecular Sciences 19, 3861 (2018).

26 Amritkar, K., Cuevas-Zuviría, B. & Kaçar, B. Evolutionary Dynamics of RuBisCO: Emergence of the Small Subunit and its Impact Through Time. Molecular Biology and Evolution 42 (2024).

27 Moreira, Sofia M., Chyou, T.-y., Wade, Joseph T. & Brown, Chris M. Diversification of the Rho transcription termination factor in bacteria. Nucleic Acids Research 52, 8979–8997 (2024).

28 Cuevas-Zuviría, B. et al. Structural evolution of nitrogenase enzymes over geologic time. eLife 14, RP105613 (2025).

29 Caserta, E. et al. Translation initiation factor IF2 interacts with the 30 S ribosomal subunit via two separate binding sites. J Mol Biol 362, 787–799 (2006).

30 Dongre, R., Folkers, G. E., Gualerzi, C. O., Boelens, R. & Wienk, H. A model for the interaction of the G3-subdomain of *Geobacillus stearothermophilus* IF2 with the 30S ribosomal subunit. Protein Sci 25, 1722–1733 (2016).

31 Wienk, H. et al. Solution structure of the C1-subdomain of *Bacillus stearothermophilus* translation initiation factor IF2. Protein Science 14, 2461–2468 (2005).

32 Guenneugues, M. et al. Mapping the fMet-tRNA(f)(Met) binding site of initiation factor IF2. Embo j 19, 5233–5240 (2000).

33 Fer, E., McGrath, K. M., Guy, L., Hockenberry, A. J. & Kaçar, B. Early divergence of translation initiation and elongation factors. Protein Sci 31, e4393 (2022).

34 Allen, G. S., Zavialov, A., Gursky, R., Ehrenberg, M. & Frank, J. The Cryo-EM Structure of a Translation Initiation Complex from *Escherichia coli*. Cell 121, 703–712 (2005).

35 Eiler, D., Lin, J., Simonetti, A., Klaholz, B. P. & Steitz, T. A. Initiation factor 2 crystal structure reveals a different domain organization from eukaryotic initiation factor 5B and mechanism among translational GTPases. Proceedings of the National Academy of Sciences 110, 15662–15667 (2013).

36 Sprink, T. et al. Structures of ribosome-bound initiation factor 2 reveal the mechanism of subunit association. Sci Adv 2, e1501502 (2016).

37 Tompa, P. Intrinsically unstructured proteins. Trends in biochemical sciences 27, 527–533 (2002).

38 Vymětal, J., Vondrášek, J. & Hlouchová, K. Sequence Versus Composition: What Prescribes IDP Biophysical Properties? Entropy 21, 654 (2019).

39 Martin, E. W. & Holehouse, A. S. Intrinsically disordered protein regions and phase separation: sequence determinants of assembly or lack thereof. Emerging Topics in Life Sciences 4, 307–329 (2020).

40 Borcherds, W., Bremer, A., Borgia, M. B. & Mittag, T. How do intrinsically disordered protein regions encode a driving force for liquid–liquid phase separation? Current Opinion in Structural Biology 67, 41–50 (2021).

41 Chiu, S.-H., Ho, W.-L., Sun, Y.-C., Kuo, J.-C. & Huang, J.-r. Phase separation driven by interchangeable properties in the intrinsically disordered regions of protein paralogs. Communications Biology 5, 400 (2022).

42 Sauer, D. B. & Wang, D.-N. Predicting the optimal growth temperatures of prokaryotes using only genome derived features. Bioinformatics 35, 3224–3231 (2019).

43 Reimer, L. C. et al. BacDive in 2022: the knowledge base for standardized bacterial and archaeal data. Nucleic Acids Research 50, D741–D746 (2021).

44 Tung Ho, L. s. & Ané, C. A Linear-Time Algorithm for Gaussian and Non-Gaussian Trait Evolution Models. Systematic Biology 63, 397–408 (2014).

45 Jabłońska, J. & Tawfik, D. S. The number and type of oxygen-utilizing enzymes indicates aerobic vs. anaerobic phenotype. Free Radical Biology and Medicine 140, 84–92 (2019).

46 Barrio-Hernandez, I. et al. Clustering predicted structures at the scale of the known protein universe. Nature 622, 637–645 (2023).

47 van Kempen, M. et al. Fast and accurate protein structure search with Foldseek. Nature Biotechnology 42, 243–246 (2024).

48 Metelev, M. & Johansson, M. A complex between IF2 and NusA suggests early coupling of transcription-translation. Nat Commun 16, 6906 (2025).

49 Gusarov, I. & Nudler, E. Control of intrinsic transcription termination by N and NusA: the basic mechanisms. Cell 107, 437–449 (2001).

50 Vvedenskaya, I. O. et al. Massively Systematic Transcript End Readout, “MASTER“: Transcription Start Site Selection, Transcriptional Slippage, and Transcript Yields. Mol Cell 60, 953–965 (2015).

51 Krupp, F. et al. Structural Basis for the Action of an All-Purpose Transcription Anti-termination Factor. Mol Cell 74, 143–157.e145 (2019).

52 Brandi, A. et al. Translation initiation factor IF2 contributes to ribosome assembly and maturation during cold adaptation. Nucleic Acids Res 47, 4652–4662 (2019).

53 Vachon, G., Raingeaud, J., Dérijard, B., Julien, R. & Cenatiempo, Y. Domain of *E. coli* translational initiation factor IF2 homologous to lambda cI repressor and displaying DNA binding activity. FEBS Lett 321, 241–246 (1993).

54 Larigauderie, G. et al. Mutation of Thr445 and Ile500 of initiation factor 2 G-domain affects *Escherichia coli* growth rate at low temperature. Biochimie 82, 1091–1098 (2000).

55 Mallikarjun, J. & Gowrishankar, J. Essential Role for an Isoform of *Escherichia coli* Translation Initiation Factor IF2 in Repair of Two-Ended DNA Double-Strand Breaks. J Bacteriol 204, e0057121 (2022).

56 Hyman, A. A., Weber, C. A. & Jülicher, F. Liquid-Liquid Phase Separation in Biology. Annual Review of Cell and Developmental Biology 30, 39–58 (2014).

57 Gao, Z. et al. Liquid-Liquid Phase Separation: Unraveling the Enigma of Biomolecular Condensates in Microbial Cells. Front Microbiol 12, 751880 (2021).

58 Moses, D., Ginell, G. M., Holehouse, A. S. & Sukenik, S. Intrinsically disordered regions are poised to act as sensors of cellular chemistry. Trends Biochem Sci 48, 1019–1034 (2023).

59 Hockenberry, A. J., Stern, A. J., Amaral, L. A. N. & Jewett, M. C. Diversity of Translation Initiation Mechanisms across Bacterial Species Is Driven by Environmental Conditions and Growth Demands. Molecular Biology and Evolution 35, 582–592 (2017).

60 Melnikov, S., Manakongtreecheep, K. & Söll, D. Revising the Structural Diversity of Ribosomal Proteins Across the Three Domains of Life. Mol Biol Evol 35, 1588–1598 (2018).

61 Ghosh, A. et al. Bacterial IF2’s N-terminal IDR drives cold-induced phase separation and promotes fitness during cold stress. bioRxiv, 2025.2002.2002.631968 (2025).

62 Libert, F., Vassart, G. & Parmentier, M. Current developments in G-protein-coupled receptors. Curr Opin Cell Biol 3, 218–223 (1991).

63 Parra-Rivas, L. A., Palfreyman, M. T., Vu, T. N. & Jorgensen, E. M. Interspecies complementation identifies a pathway to assemble SNAREs. iScience 25, 104506 (2022).

64 Shimasaki, N. B. & Kane, C. M. Structural basis for the species-specific activity of TFIIS. J Biol Chem 275, 36541–36549 (2000).

65 Hershey, J. W. B., Sonenberg, N. & Mathews, M. B. Principles of Translational Control. Cold Spring Harb Perspect Biol 11 (2019).

66 Lynch, M. & Marinov, G. K. The bioenergetic costs of a gene. Proc Natl Acad Sci U S A 112, 15690–15695 (2015).

67 Parks, D. H. et al. GTDB: an ongoing census of bacterial and archaeal diversity through a phylogenetically consistent, rank normalized and complete genome-based taxonomy. Nucleic Acids Research 50, D785–D794 (2021).

68 Nadkarni, P. M. & Parikh, C. R. An eUtils toolset and its use for creating a pipeline to link genomics and proteomics analyses to domain-specific biomedical literature. J Clin Bioinforma 2, 9 (2012).

69 Katoh, K. & Standley, D. M. MAFFT Multiple Sequence Alignment Software Version 7: Improvements in Performance and Usability. Molecular Biology and Evolution 30, 772–780 (2013).

70 Nguyen, L.-T., Schmidt, H. A., von Haeseler, A. & Minh, B. Q. IQ-TREE: A Fast and Effective Stochastic Algorithm for Estimating Maximum-Likelihood Phylogenies. Molecular Biology and Evolution 32, 268–274 (2014).

71 Kalyaanamoorthy, S., Minh, B. Q., Wong, T. K. F., von Haeseler, A. & Jermiin, L. S. ModelFinder: fast model selection for accurate phylogenetic estimates. Nature Methods 14, 587–589 (2017).

72 Tria, F. D. K., Landan, G. & Dagan, T. Phylogenetic rooting using minimal ancestor deviation. Nat Ecol Evol 1, 193 (2017).

73 Mirdita, M. et al. ColabFold: making protein folding accessible to all. Nat Methods 19, 679–682 (2022).

74 Suyama, M., Torrents, D. & Bork, P. PAL2NAL: robust conversion of protein sequence alignments into the corresponding codon alignments. Nucleic Acids Res 34, W609–612 (2006).

75 Yang, Z. PAML 4: Phylogenetic Analysis by Maximum Likelihood. Molecular Biology and Evolution 24, 1586–1591 (2007).

76 Emenecker, R. J., Griffith, D. & Holehouse, A. S. Metapredict: a fast, accurate, and easy-to-use predictor of consensus disorder and structure. Biophys J 120, 4312–4319 (2021).

77 Hatos, A., Tosatto, S. C. E., Vendruscolo, M. & Fuxreiter, M. FuzDrop on AlphaFold: visualizing the sequence-dependent propensity of liquid-liquid phase separation and aggregation of proteins. Nucleic Acids Res 50, W337–w344 (2022).

78 Winter, D. J. rentrez: An R package for the NCBI eUtils API. Report No. 2167-9843, (PeerJ Preprints, 2017).

