## Supplementary Information for "Evolution of Translation Initiation Factor 2 Extensions Links Initiation to Bacterial Stress Response"

**Extended Data 1:** The list of IF2 types

**Extended Data 2:** The list of organisms used in the dataset with their IF2 N- and C-terminal extension lengths, predicted optimal growth temperatures, and oxygen preference, NusA protein accession list and NusA C-terminal extension lengths

**Extended Data 3:** The list of strains, plasmid constructs, and primer sequences used in this study

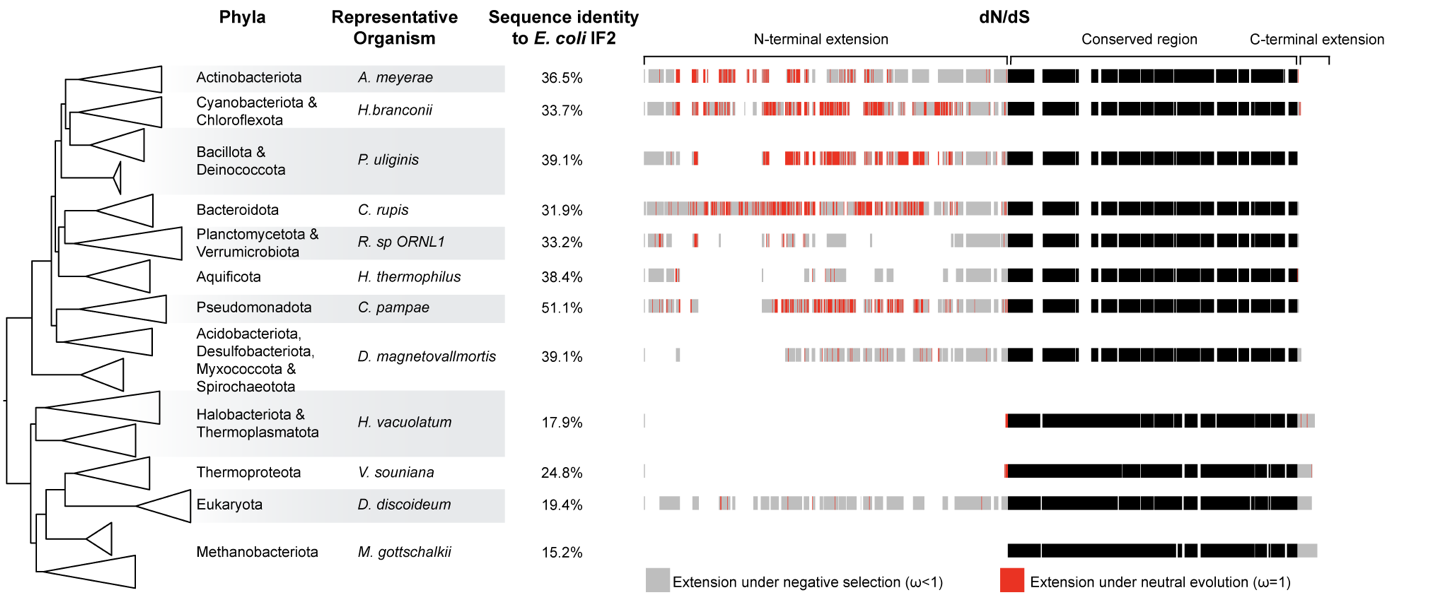

**Extended Data Fig 1.** The simplified tree representing bacterial, archaeal and eukaryotic phyla groups with a representative organism to show dN/dS (*ω*) values on the sequence. The selection analysis is projected onto the alignment of representative sequences. Gray indicates strong purifying selection (*ω* < 1), whereas red denotes sites under neutral evolution (*ω*=1). Black regions indicate the conserved internal regions across IF2 proteins. *A. meyerae: Actinomadura meyerae*; *H. branconii*: *Halotia branconii*; *P. uliginis*: *Paenibacillus uliginis*; *C. rupis*: *Chitinophaga rupis*; *R. sp ORNL1*: *Roseimicrobium sp. ORLN1*; *H. thermophilus*: *Hydrogenobacter thermophilus*; *C. pampae*: *Cupriavidus pampae*; *D. magnetovalmortis*: *Desulfamplus magnetovalmortis*; *H. vacuolatum*: *Halorubrum vacuolatum*; *V. souniana*: *Vulcanisaeta souniana*; *D. discoideum*: *Dictyostelium discoideum*; *M. gottschalkii*: *Methanobrevibacter gottschalkii.*

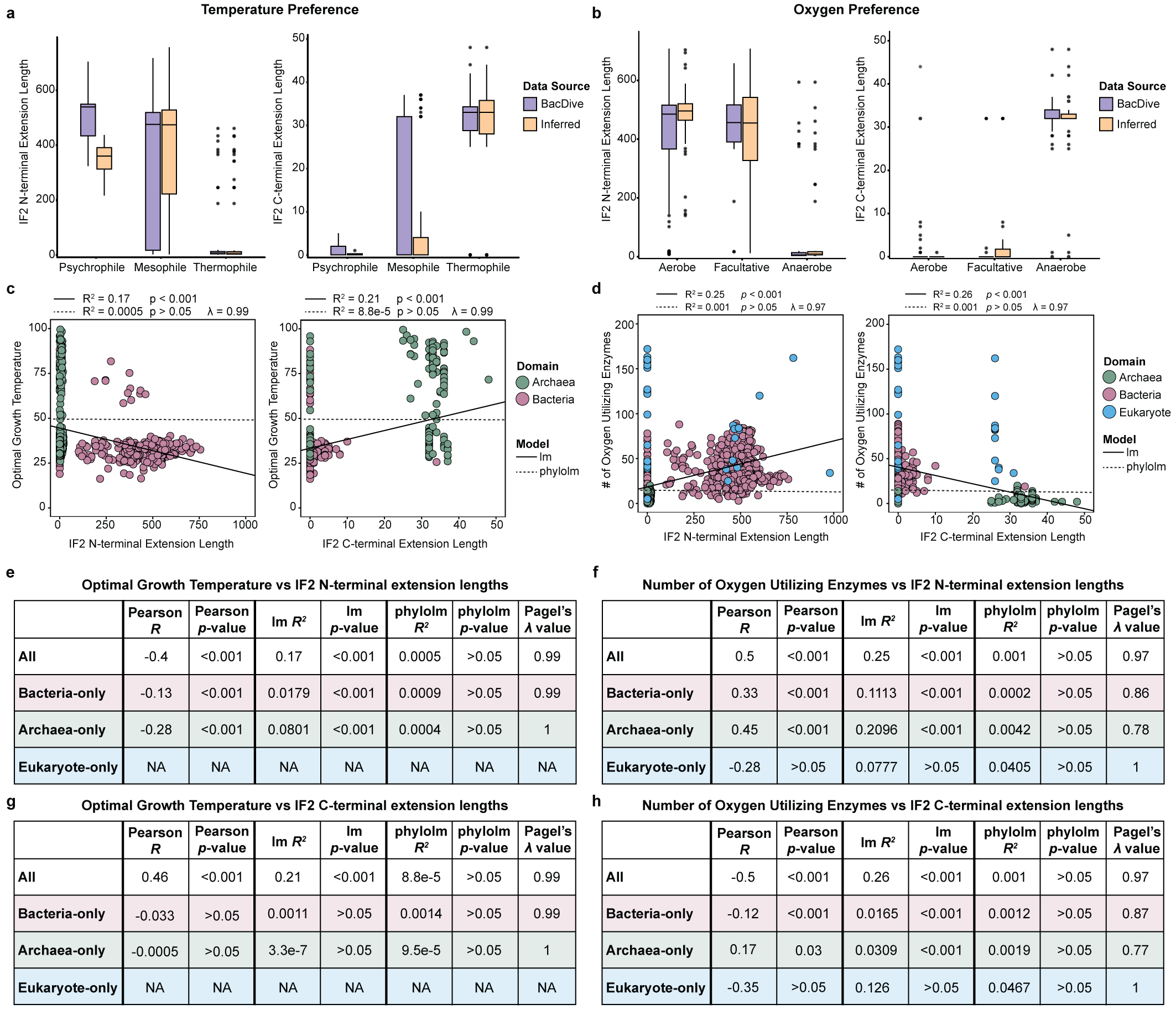

**Extended Data Fig 2.** Correlations between IF2 extension lengths and environmental preferences. **a)** Boxplots showing the distribution of IF2 N-terminal (left) and C-terminal (right) extension lengths across organisms categorized by temperature preference: psychrophiles, mesophiles, and thermophiles. **b)** Boxplots showing the distribution of IF2 N-terminal (left) and C-terminal (right) extension lengths across organisms with different oxygen preferences: aerobes, facultative, and anaerobes. Data are sourced either from BacDive (purple) or inferred based on genome information (yellow). **c)** Correlations between optimal growth temperature and IF2 N-terminal (left) and C-terminal (right) extension lengths. Solid lines represent linear regression (lm), and dashed lines represent phylogenetic regression with *λ* model (phylolm). Points are colored by domain (Bacteria: pink, Archaea: green). **d)** Correlations between the number of oxygen-utilizing enzymes and IF2 N-terminal (left) and C-terminal (right) extension lengths. Solid and dashed lines represent lm and phylolm models, respectively. Points are colored by domain (Bacteria: pink, Archaea: green, Eukaryotes: blue).

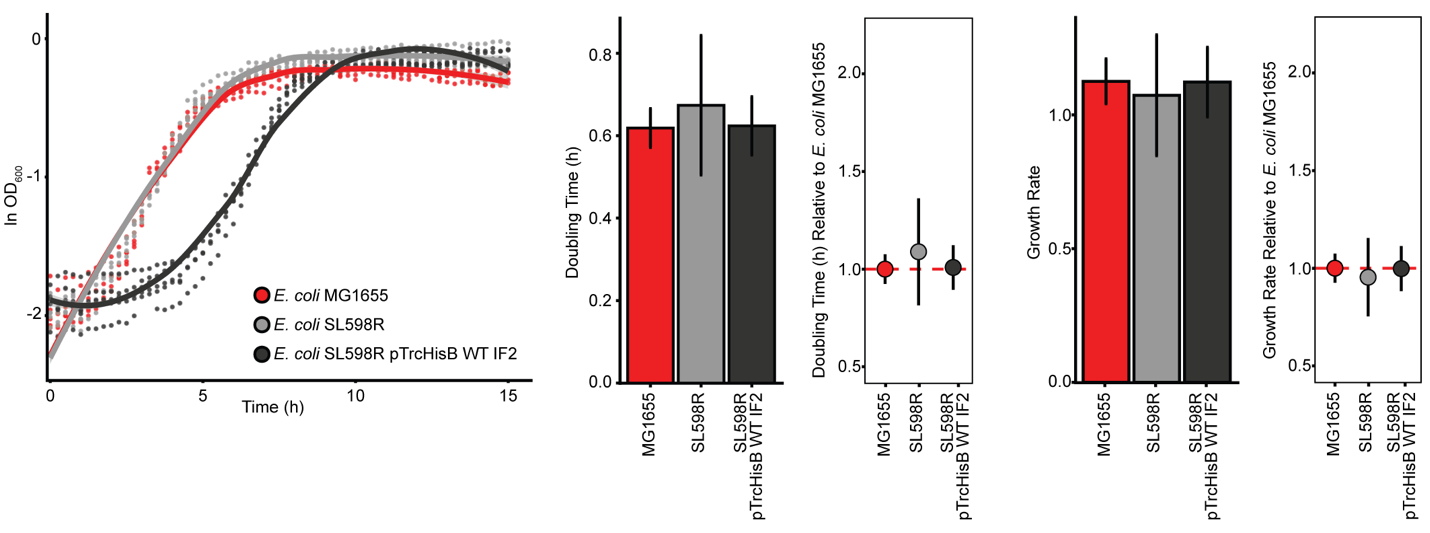

**Extended Data Fig 3.** Growth comparison between different *E. coli* strains: *E. coli* MG1655; *E. coli* SL598R; *E. coli* SL598R with pTrcHisB-WTIF2 construct (heat curated for 30 min and checked for removal of endogenous IF2 from the genome). Doubling time and growth rates are shown as barplots. The doubling time and growth rates relative to *E. coli* MG1655 are shown as boxplots. None of the comparisons gave significant difference for growth based on Wilcoxon rank-sum test (*p*-value>0.05).

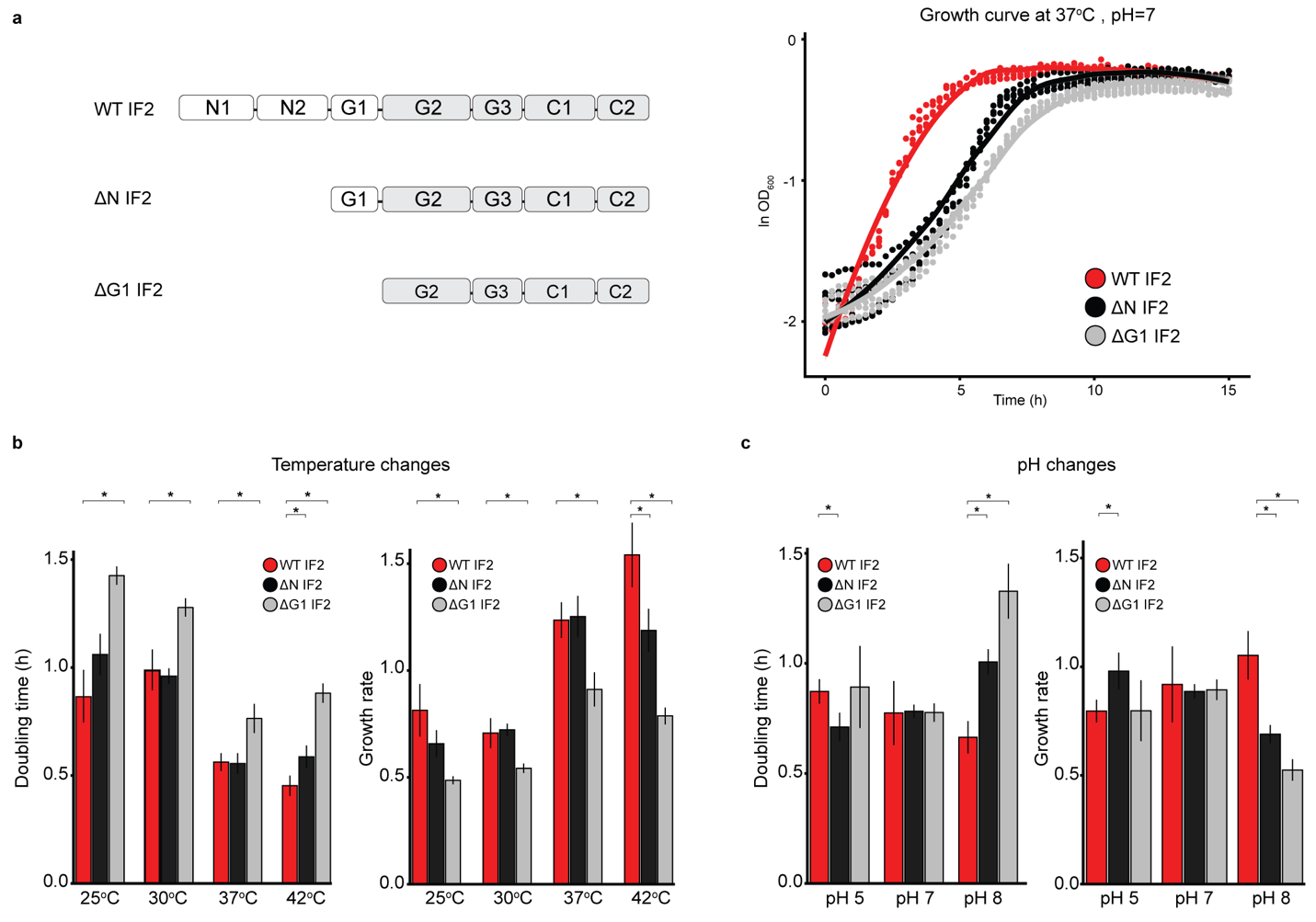

**Extended Data Fig 4.** Growth with IF2 variants at different temperature and pH. **a)** The overview of the IF2 variants used in the study. The growth curves of *E. coli* with different IF2s at 37^o^C and pH 7 (right). **b)** Growth rate and doubling times at various temperatures (25°C, 30°C, 37°C, 42°C). **c)** Growth rate and doubling times at various pH (pH 5, pH 7, pH 8). *E. coli* with wild-type IF2 (WT IF2) is shown as red; N1-N2 domain truncated IF2 (ΔN IF2) as black; and N1-N2-G1 domain-truncated IF2 (ΔG1 IF2) as gray. Each barplots represents the doubling times and growth rates of five replicates (n=5). Asterisks indicate statistically significant differences by Wilcoxon signed-rank test (*p*-value<0.05).

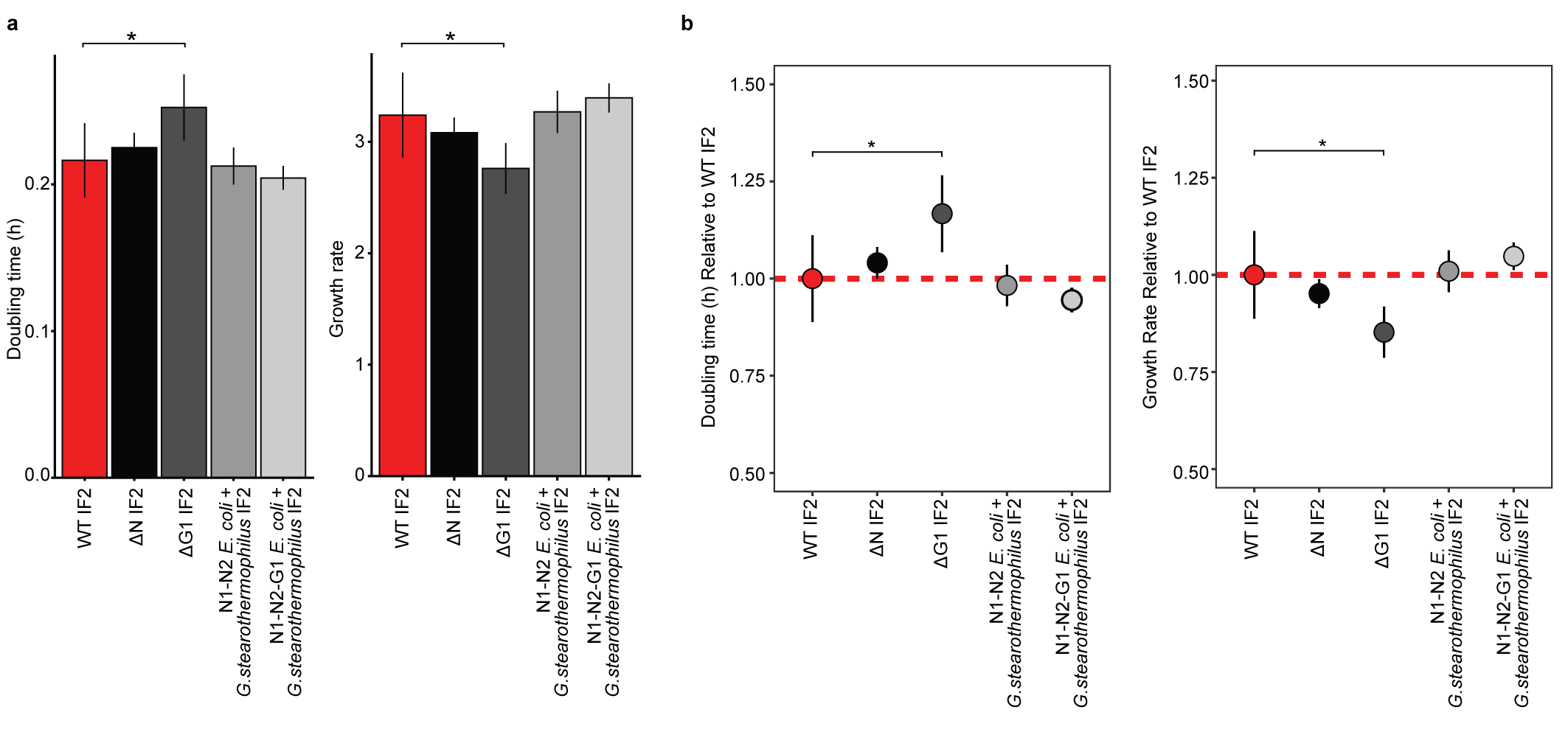

**Extended Data Fig 5.** Doubling time and growth rate comparison for N-terminal swaps between *G. stearothermophilus* and *E. coli* IF2. **a)** Bar plots showing doubling times and growth rates of IF2 variants, including wild-type IF2 (*E. coli* IF2), N-terminal truncations (ΔN and ΔG1), and chimeric constructs in which *E. coli* N1–N2 or N1–N2–G1 domains are fused to the *G. stearothermophilus* IF2 core. **b)** Box plots showing doubling times and growth rates of IF2 variants normalized to wild-type IF2 (*E. coli* IF2). Each condition includes eight biological replicates. Asterisks denote statistically significant differences relative to wild type as determined by a Wilcoxon rank-sum test (p-value<0.05). The red dashed line indicates the wild-type IF2 reference value (doubling time = 1).

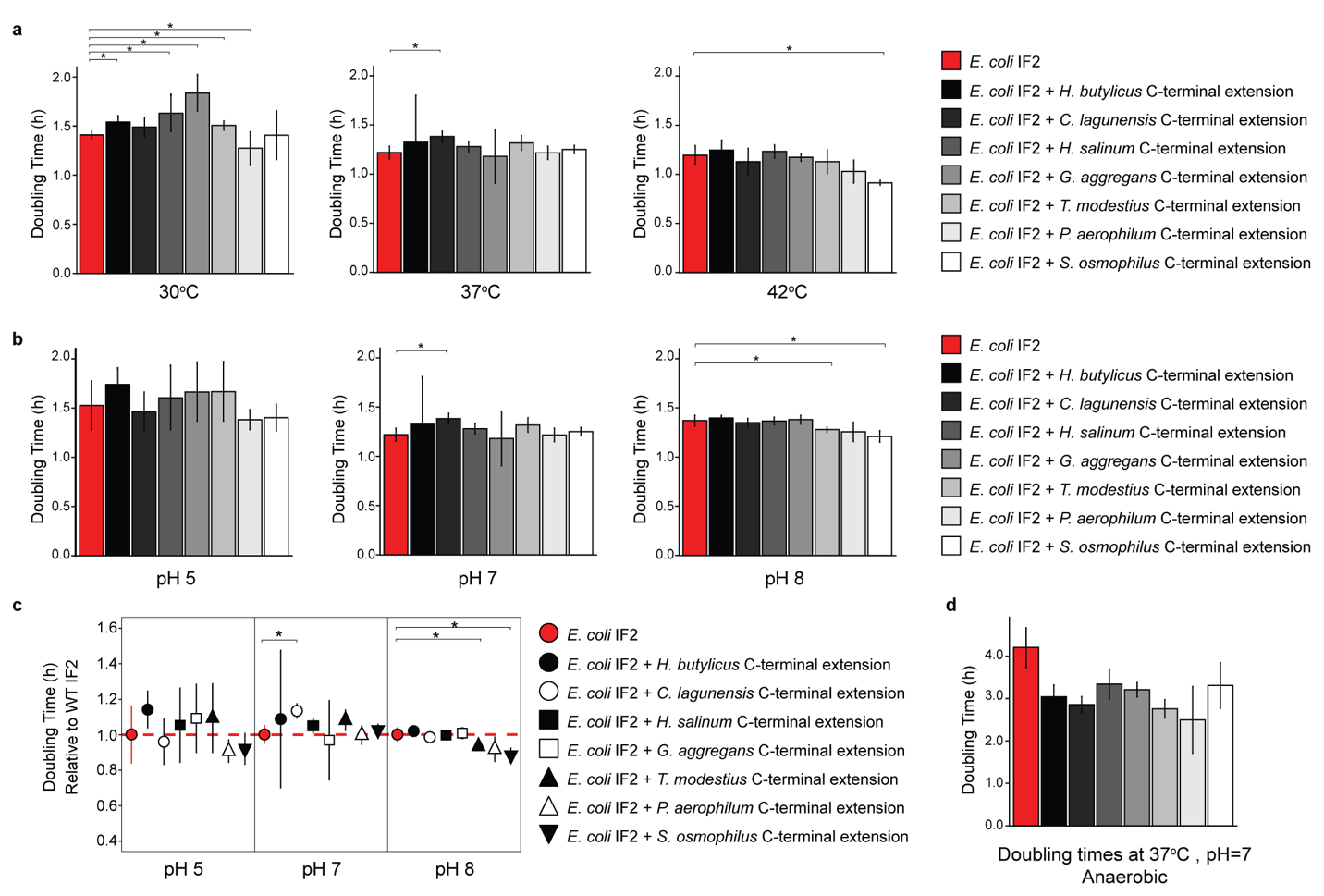

**Extended Data Fig 6.** Barplots of doubling time at various **a)** temperatures (30^o^C, 37^o^C, 42^o^C) and **b)** pH (pH 5, pH7, pH8) are shown relative to the WT IF2. **c)** Boxplots of doubling times relative to WT IF2 at various pH. **d)** Barplots of doubling time at 37^o^C, pH 7 and anaerobic condition. Each point represents the distribution of relative doubling time of five replicates (n=5). Asterisks indicate statistically significant differences by Wilcoxon rank-sum test (*p*-value<0.05). The red dashed line indicates WT IF2 doubling time at 1 as reference. *H. butylicus*: *Hyperthermus butylicus*; *C. lagunensis*: *Caldisphaera lagunensis*; *H. salinum*: *Halorarum salinum*; *G. aggregans*: *Granulicella aggregans*; *T. modestius*: *Thermocladium modestius*; *P. aerophilum*: *Pyrobaculum aerophilum*; *S. osmophilus*: *Schizosaccharomyces osmophilus*.

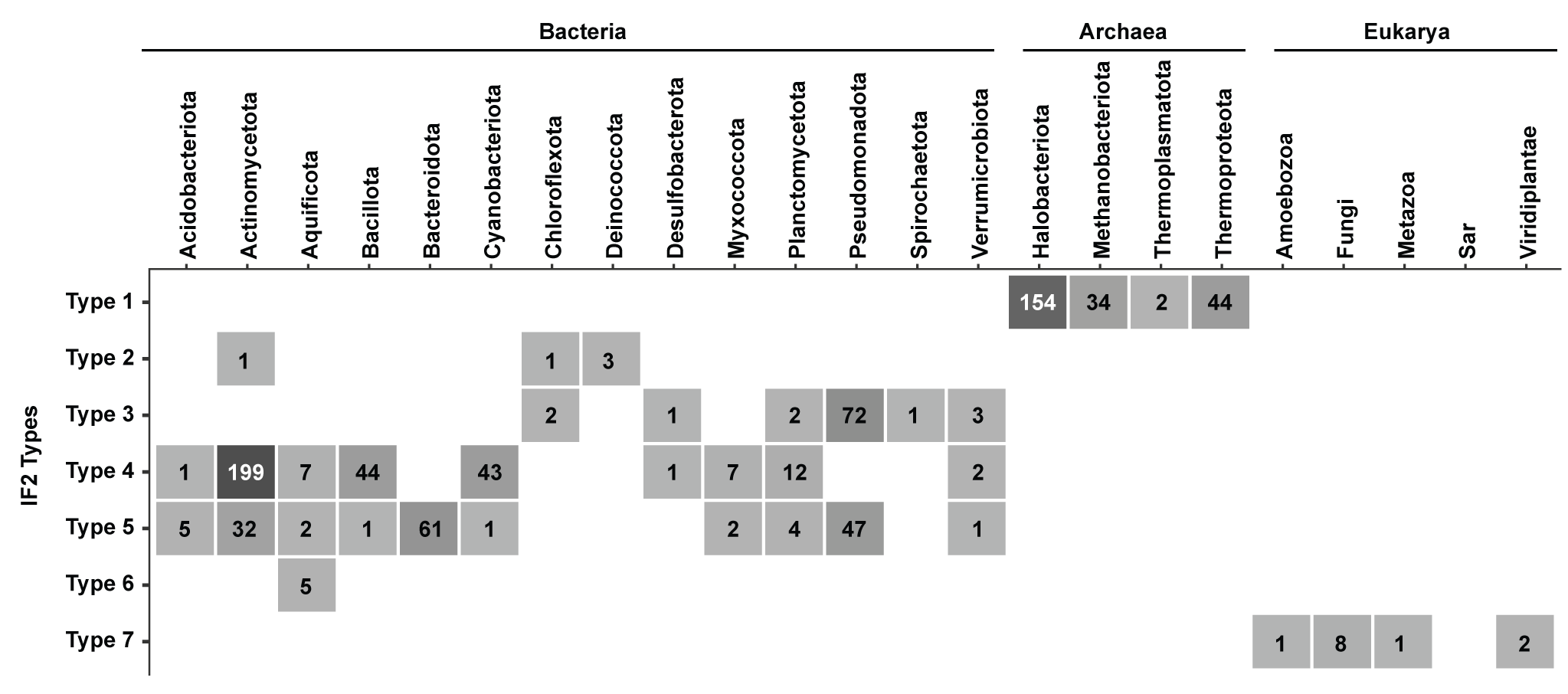

**Extended Data Fig 7.** Distribution of IF2 types across major bacterial, archaeal, and eukaryotic clades.

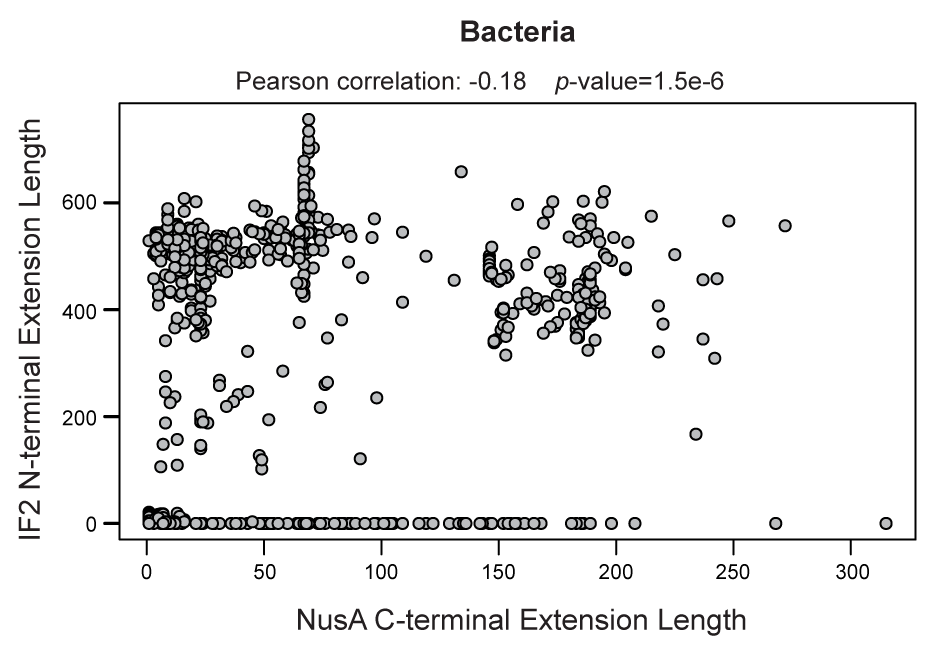

**Extended Data Fig 8**. A scatter plot showing the relationship between NusA C-terminal extension length and IF2 N-terminal extension length across bacterial organisms. Each point represents a distinct bacterial species.

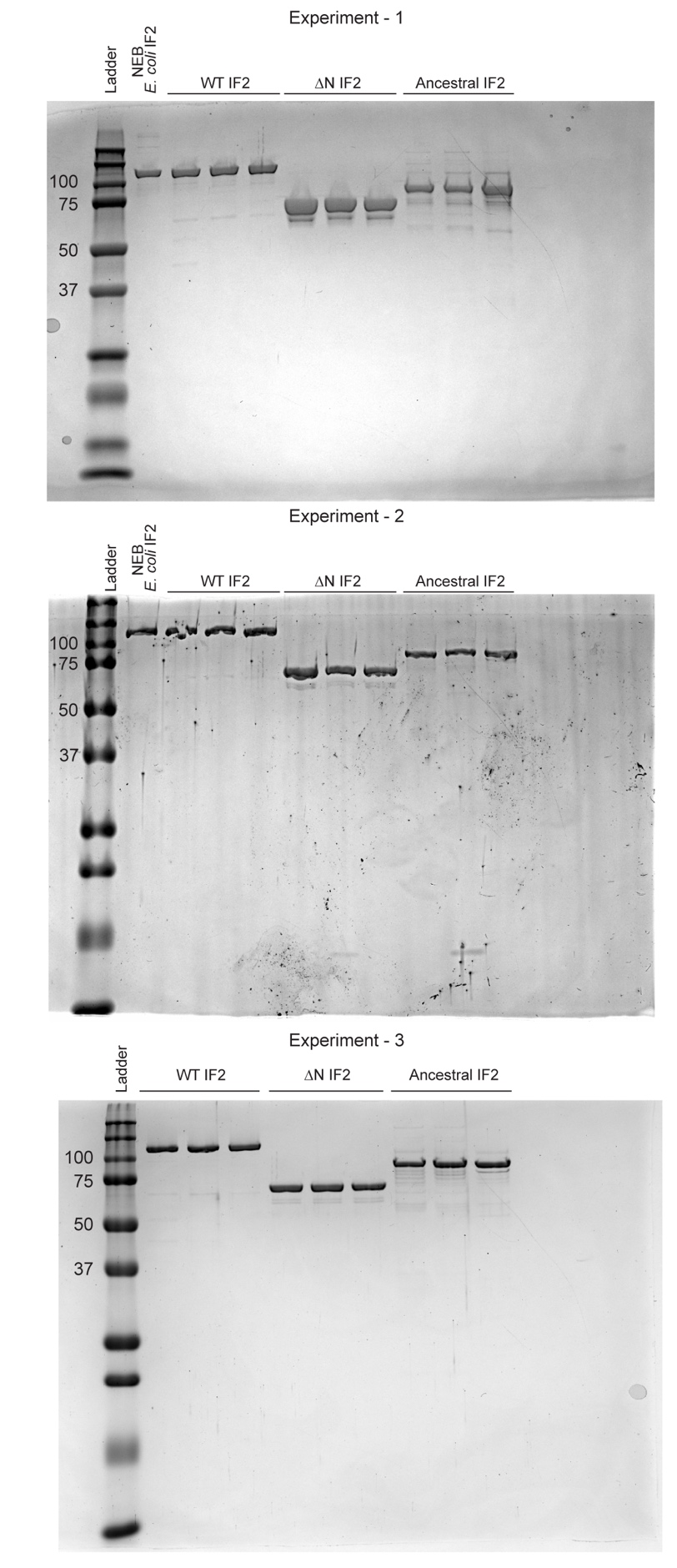

**Extended Data Fig 9**. SDS-PAGES for the normalized concentrations of purified IF2 variants before *in vitro* experiment. In each experiment, 3 replicates of purified IF2 variants were used. For experiment 1, the purified IF2 variants were normalized to 3 ug of purified WT IF2. For experiment 2 and 3, the purified IF2 variants were normalized to 2 ug of purified WT IF2. An ancestral IF2 was used in normalization however not included in the in vitro experiments in this study. NEB E. coli IF2: New England Biotechnology supplied *E. coli* IF2. The expected band sizes in kDa are shown for ladder on the left side of each gel.
